# Mdivi-1 affects neuronal activity by inhibition of Complex I and respiratory supercomplex assembly

**DOI:** 10.1101/2024.01.25.577160

**Authors:** Nico Marx, Nadine Ritter, Paul Disse, Guiscard Seebohm, Karin B. Busch

**Affiliations:** Institute for Integrative Cell Biology and Physiology (IIZP), Schloßplatz 5, Department of Biology, University of Münster, 48149 Münster, Germany; Institute for Genetics of Heart Diseases (IfGH), Department of Cardiovascular Medicine, University Hospital Münster, D-48149 Münster, Germany; Department of Drug Design and Pharmacology, University of Copenhagen, DK-2100 Copenhagen, Denmark

**Keywords:** Mitochondria, respiratory Complex I, Mdivi-1, inhibition, respiratory supercomplexes, neuronal activity, calcium metabolism

## Abstract

Several human diseases, including cancer and neurodegeneration, are associated with excessive mitochondrial fragmentation. In this context, mitochondrial division inhibitor (Mdivi-1) has been tested as a therapeutic to block the fission-related protein dynamin-like protein-1 (Drp1). Recent studies suggest that Mdivi-1 interferes with mitochondrial bioenergetics. Here we show that the molecular mechanism of Mdivi-1 is based on inhibition of complex I at the IQ site. This leads to the destabilization of complex I, impairs the assembly of N- and Q-respirasomes and is associated with increased ROS production. The result is a reduced efficiency of ATP generation. Second, the calcium homeostasis of cells is impaired, which severely affects the electrical activity of neurons. Given the results presented here, a potential therapeutic application of Mdivi-1 is challenging because of its impact on synaptic activity. Similar to the Complex I inhibitor rotenone, Mdivi-1 may lead to neurodegenerative effects in the long term.

**Graphical abstract:** 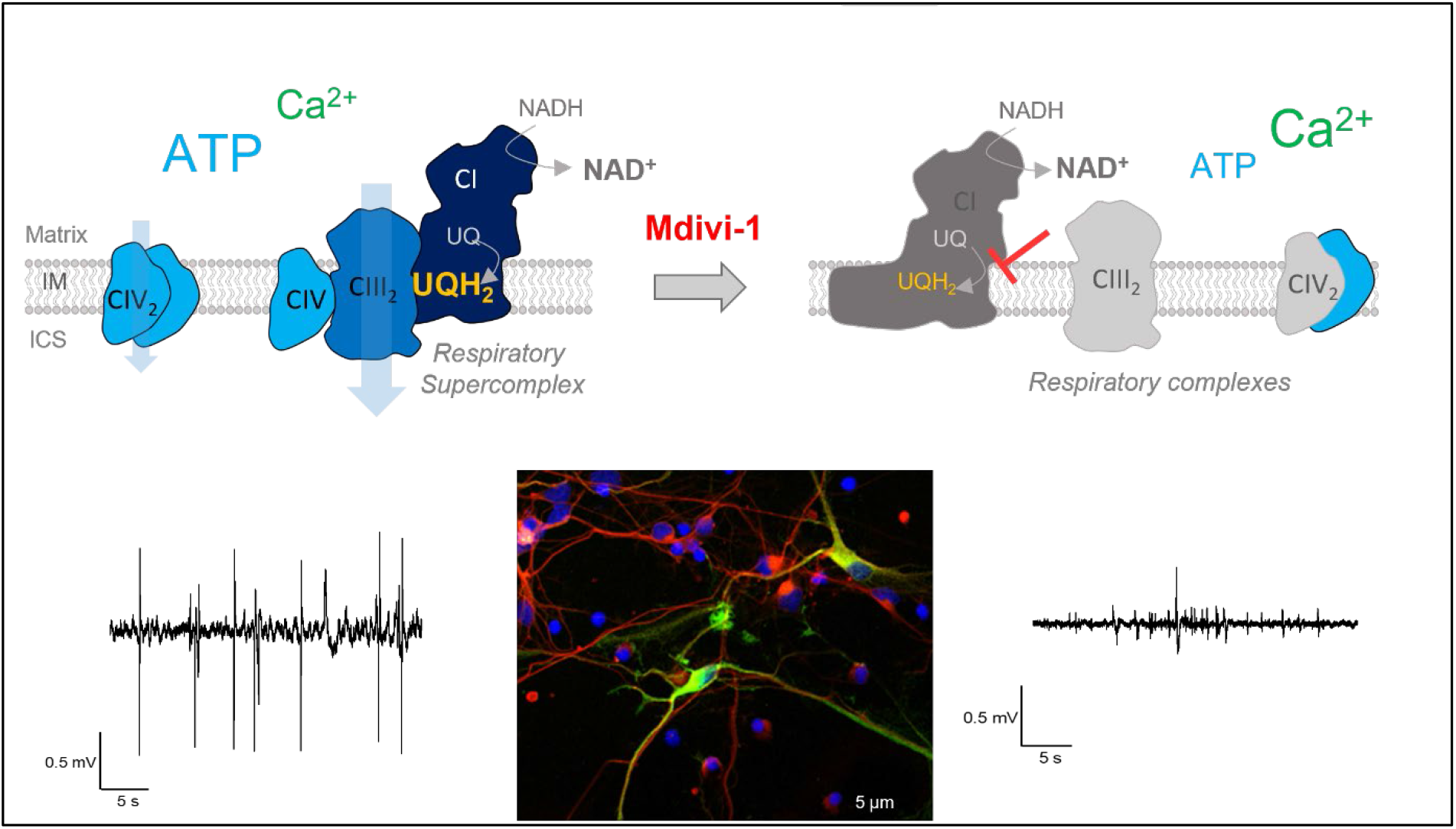

- Mdivi-1 inhibits respiratory complex I at the IQ-site
- Inhibition destabilizes complex I and reduces supercomplex formation
- Mitochondrial ATP levels decrease
- Ca^2+^ metabolism is affected
- Neuronal activity is compromised

## Introduction

Neurons rely on mitochondrial ATP synthesis to fuel energy-intensive processes such as vesicle cycling and signal transduction. Consequently, mitochondria dysfunction is closely associated with neurodegenerative diseases (Cavallucci, Nobili et al. 2013, Lee, Hirabayashi et al. 2018). In particular, respiratory Complex I (CI) inhibition seems to facilitate initiation of neurodegeneration (Greenamyre, Sherer et al. 2001, Schapira 2010). Mammalian Complex I (CI), the NADH-ubiquinone oxidoreductase, is one of the largest multiprotein membrane complexes with a molecular weight of about 1 MDa. It consists of 45 subunits. CI has three catalytic functions: NADH oxidation, electron transfer to ubiquinone, and proton translocation. Together with respiratory complexes CIII and CIV, CI generates the proton motive force across the mitochondrial inner membrane (IMM), which drives ATP synthesis by ATP synthase (CV). CII is the succinate dehydrogenase, CIII the ubiquinol-cytochrome *c* oxidoreductase and CIV the cytochrome c-oxidase.

CI catalyzes the oxidation of NADH to NAD^+^ at the tip of the peripheral stalk. The electrons are transported via cofactors from the N-module to the Q-module. N- and Q-module build the hydrophilic peripheral arm. The flow of electrons alters the redox state of the protein and induces conformational changes of the protein. NADH-induced changes in the Q-cavity are only possible in the open conformation of the complex, while the reduction of quinone happens only in the closed state (Kampjut and Sazanov 2020). The coupling mechanism between electron transfer and proton translocation in Complex I remains a controversial question in the field (Parey, Lasham et al. 2021, Chung, Wright et al. 2022, Kampjut and Sazanov 2022). The membrane-embedded part, which transports protons from the matrix into the intracristal space, consists of the P_D_- and P_P_-module. Mutations in CI subunits can disrupt the proper assembly of CI, leading to unstable CI subassemblies and dysfunction including increased formation of reactive oxygen species (ROS) (Mimaki, Wang et al. 2012, Formosa, Muellner-Wong et al. 2020). Chemical inhibition of CI can induce parkinsonism in humans. The inhibitor meperidine selectively targeted dopaminergic neurons of the *Substantia nigra* and induced dopamine deficiency and cell death (Langston, Ballard et al. 1983).

In neurons, fusion and fission dynamics of mitochondria is important for the adaptation of mitochondrial and cellular functions (Misgeld and Schwarz 2017). So, mitochondrial fission is a prerequisite for the maturation of neuronal progenitor cells (NPCs) during neurogenesis (Iwata, Casimir et al. 2020, Brunetti, Dykstra et al. 2021). Fission is mediated by the cytosolic dynamin-related protein Drp1, a GTPase. The chemical agent Mitochondrial Division Inhibitor-1 (Mdivi-1) is a commonly used inhibitor of Drp1 to suppress mitochondrial fission (Manczak, Kandimalla et al. 2019), which is also effective in primary neurons. Several studies provided evidence of protective effects of fission-inhibition by Mdivi-1 in neurodegenerative models for Alzheimer’s disease (AD) (Baek, Park et al. 2017, Reddy, Manczak et al. 2017) and Parkinson’s disease (PD) (Rappold, Cui et al. 2014, Bido, Soria et al. 2017). However, more recent studies suggested that Mdivi-1 effects are more complex and target mitochondrial bioenergetics (Bordt, Clerc et al. 2017, Zhang, Wang et al. 2017, Ruiz, Alberdi et al. 2018). Clarification of this issue is important because inhibition of the ETC, in particular Complex I, is closely associated with onset of neurodegeneration (Greenamyre, Sherer et al. 2001, Schapira 2010). Here we provide evidence that Mdivi-1 directly inhibits Complex I by binding to the I_Q_ site. This affects the assembly of Complex I and respiratory supercomplexes with consequences for ATP generation and mitochondrial calcium homeostasis. Ultimately, this leads to functional impairment of neurons.

## Results

The Mdivi-1 effect was tested in two human cell model systems: Hela cells, a cervical cancer cell line, and neurons differentiated from neuronal progenitor cells (NPC). The generation of functional NPC-derived midbrain neurons was confirmed by immune-staining and electrophysiological characterization (Supplementary Figure S1).

### Effects of Mdivi-1 on mitochondrial and cellular function

#### Mdivi-1 affects cell growth and mitochondrial biogenesis

To check general Mdivi-1 effects on cellular health, growth curves of HeLa cells were recorded. Long-term and acute treatment with Mdivi-1 as well as 24 h treatment with 1 μM rotenone significantly impaired the growth in comparison to DMSO treated control cells (Figure 1A). Mdivi-1 treatment also resulted in significantly increased p21 expression, which is a cyclin-dependent kinase (CDK1) inhibitor (Figure 1B). Cell cycle analysis by flow cytometry after propidium iodide (PI) staining allowed to dissect differences in relative cell cycle stages. Treatment of HeLa cells with Mdivi-1, rotenone or nocodazole, an inhibitor of microtubule dynamics and inducer of cell cycle arrest, led to a shift from G0/G1 phase to S-Phase (Supplementary Figure S2). Nocodazole (24h, 200 nM) furthermore induced a significant increase of cells in G2/M phase. NPC, which were acutely treated with Mdivi-1 showed an increase in S and G2/M phase (Supplementary Figure S2). These shifts in cell cycle states from G0/G1 to S and G2/M phase suggest an impaired cellular growth due to stochastic failure of mitosis. Mdivi-1 treatment also affected cell morphology: NPC displayed a morphological change (Figure 1C) and cells became rounder (Figure 1D).

**Figure 1:**
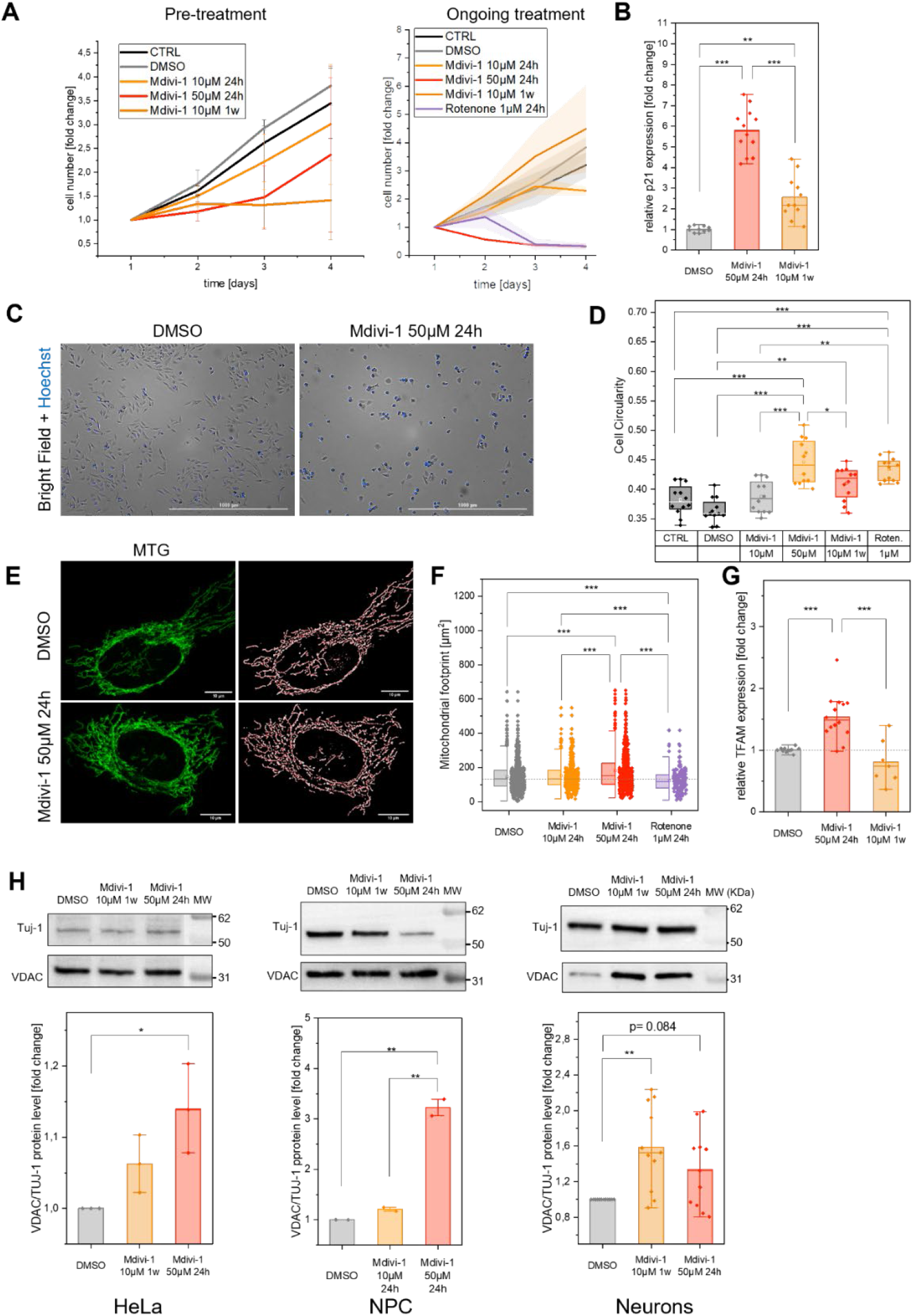
Mdivi-1 affects cell growth and mitochondrial biogenesis. (A) Growth curves of cells treated with Mdivi-1 for 24h, respectively 1 w, before recording. Left: pre-treatment with Mdivi-1, no Mdivi-1 during recording; right: ongoing treatment with Mdivi-1 (right). (B) Expression of p21 is elevated in Mdivi-1 treated HeLa cells (N=4, n=16, ANOVA. (C) Midivi-1 effects on cellular morphology of HeLa cells stained with Hoechst (scale bar: 1000 µm). (D) Circularity of HeLa cells with acute and long-term Mdivi-1 and Rotenone treatment (N=4 independent experiments, n=13, ANOVA). (E) Exemplary cLSM images and MiNA skeleton of control and Mdivi-1-treated HeLa cells stained with Mito Tracker™Green (MTG) (scale bar: 10 ^µ^m). (F) Treatment with Mdivi-1 (50 µM) leads to increased, while Rotenone (1 µM) leads to decreased mitochondrial mass (N=6; n_DMSO_=828, n_Mdivi-1 10 µM 24h_=519, n_Mdivi-1 50 µM 24h_=522, n_Drp1 K38A mt EGFP_=28, n_Rotenone_=196; DMSO median as dashed line). (G) Accute Mdivi-1 treatment indicates elevation in gene expression of TFAM in HeLa cells (N=4, n=16). (H) Protein levels of Voltage Dependent Anion channel (VDAC) normalized on *β*-III-Tubulin (TUJ-1) are increased in Mdivi-1-treated cells (left panel: HeLa n=3, NPC, n=2 right panel:neurons n=11, all ANOVA). Boxplots indicate median (line), 25th-75th percent percentile (box) and minimum and maximum values (whiskers). Statistics: ***p≤0,001; **p≤0,01; **p≤0,05.

To check for Mdivi-1 effects on mitochondria, treated HeLa cells were stained with MitoTracker™Green, imaged and then analyzed to determine the mitochondrial mass per cell (Figure 1E, F). Cells treated with 50 µM Mdivi-1 for 24 h had a significantly increased mitochondrial footprint. In parallel, the expression of TFAM, the mitochondrial transcription factor A, was increased indicating biogenesis of mitochondria (Figure 1G). Finally, we determined VDAC protein levels in three Mdivi-1 treated cell types (Hela, NPC and neurons). VDAC protein, normalized on Tuj-1, a class III beta-tubulin, was higher in Hela and NPC that were treated with Mdivi-1. Neurons showed a tendency towards higher VDAC-protein levels. Together, this data indicates that Mdivi-1 affects cell proliferation and cellular shape and evokes mitochondrial biogenesis.

#### Mdivi-1 treatment causes decline of mitochondrial function and increases ROS levels

To test, whether Mdivi-1 affects mitochondrial function, oxygen consumption rates (OCR) were measured with a real-time kinetic assay (Seahorse XF96 Analyzer/Agilent) (Figure 2A). Basal, ATP-synthesis-linked and maximal respiration were determined in control and Mdivi-1-treated cells before and after addition of the inhibitor oligomycin and the uncoupler FCCP, respectively. In the last step, complex I and complex III were inhibited by addition of rotenone and antimycin A (AA). OCR was normalized on the cell number. In Hela cells (Figure 2) and neurons (Supplementary Figure S3), the OCR was not significantly altered by a 24h Mdivi-1 treatment (10 and 50 µM). The metabolic profile, i.e. the ratio of the OCR and the extracellular acidification rate (ECAR), of cells treated with 50 µM Mdivi-1 for 24 h showed a tendency towards a compromised oxidative energy metabolism, while glycolysis increased (Figure 2B). However, when OCR was normalized on the mitochondrial mass per cell (which was increased in Mdivi-1-treated cells), basal, ATP synthesis-linked and maximal respiration in both, Hela cells (Figure 2C) and neurons (Figure S3) were significantly decreased compared to the control. The OCR/ECAR ratio was significantly decreased in Mdivi-1-treated cells when normalized on the number of mitochondria per cell (Figure 2D). This suggests that the observed biogenesis of mitochondria (Figure 1G) as a stress response could not fully compensate the decline of mitochondrial functionality. To further elucidate the consequences of decreased OCR, mitochondrial ATP levels were determined by using the fluorogenic dye ATPRed-1m, which fluorescence intensity is ATP-dependent. To normalize on mitochondrial mass, cells were in addition stained with MTG (Figure 2E). Indeed, relative mitochondrial ATP levels were decreased in Mdivi-1 treated cells. Remarkably, the effect was stronger than for the well characterized complex I inhibitor rotenone (Figure 2F).

**Figure 2:**
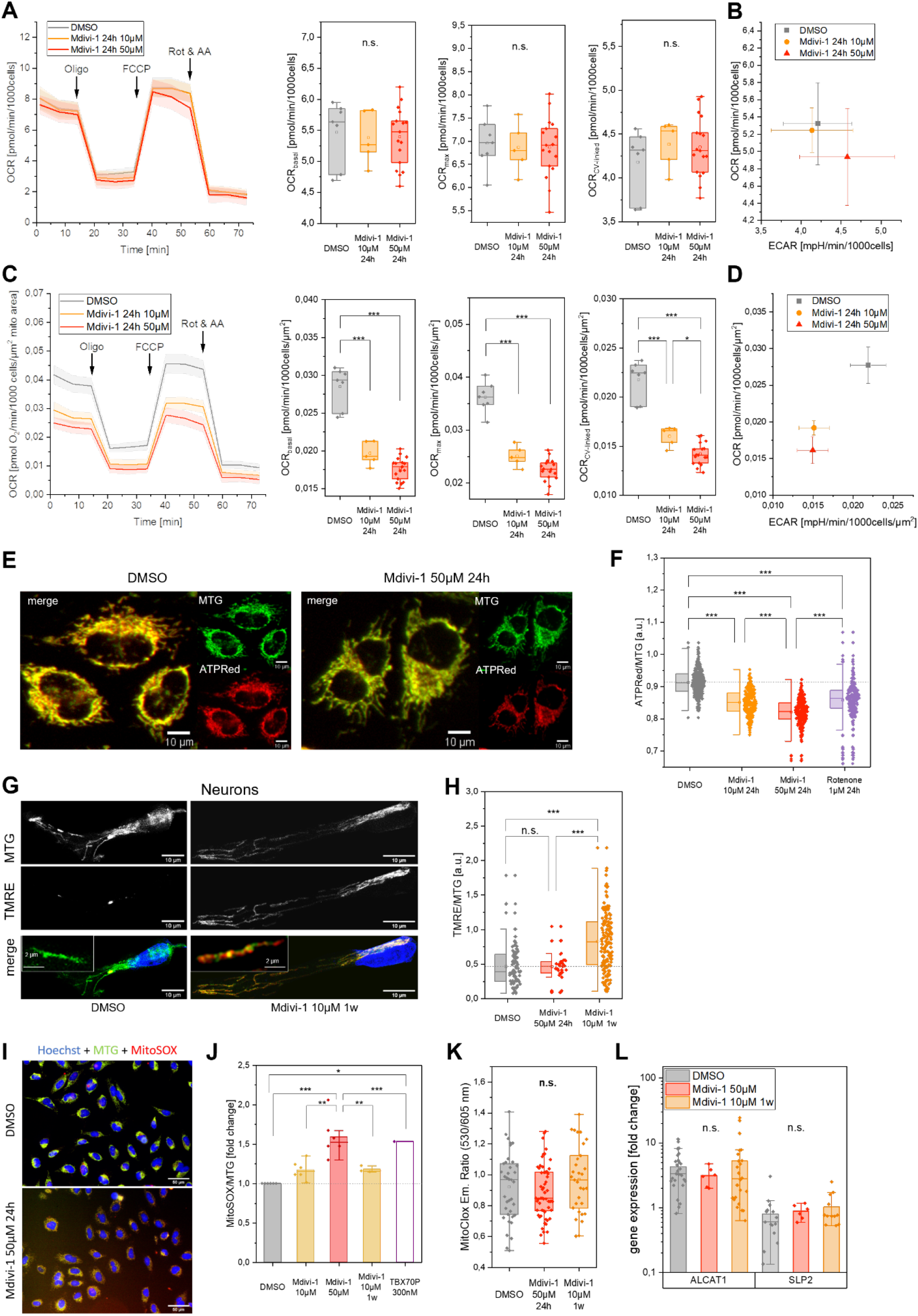
Mitochondrial function is reduced in Mdivi-1 treated cells. (A) Functional test of mitochondrial respiration with subsequent addition of an inhibitor for ATP synthase (oligomycin,1.5 μM), uncoupler trifluoromethoxy carbonylcyanide phenylhydrazone (FCCP, 2 μM) and inhibitors for complex I (Rot., Rotenone; 0.5 μM) and complex III (AA; antimycin A; 1 μM). Oxygen consumption rates (OCR) of control and Mdivi-1-treated HeLa cells were determined with an automatic flux analyzer (Seahorse XF96/Agilent) and normalized to cell numbers. (B) Metabolic profile as OCR/ECAR ratio. (C) OCR normalized on the mean mitochondrial footprint (mitochondrial area in µm^2^ per cell) determined by MiNA (Figure S1). Basal, maximal and ATP production related respiration in living cells (control and Mdivi-1 treated) were determined (N=1; n=8,5,17). (D) Metabolic profile normalized on mitochondrial area per cell. (E) Visualization of mitochondrial ATP levels in Mdivi-1 treated and control HeLa cells, co-staining with Mito Tracker Green and ATPRed-1. (F) Mitochondrial ATP is decreased in Mdivi-1- and Rotenone-treated HeLa cells (N=3, n_DMSO_=305, n_Mdivi-1/10 μM 24h_=200, n_Mdivi-1 /50 μM 1w_=235, n_Rotenone 1 μM 24h=_234). (G) Staining of mitochondrial membrane potential in Mdivi-1 treated neurons with TMRE; MTG for mitochondrial mass determination. Insets showing magnified views. (H) Mitochondrial membrane potential in neurons (N=2, n_DMSO_=87, n_Mdivi-1 10 μM 1w_=154, n_Mdivi-1 50 μM 24h_=35). (I) Exemplary images of HeLa cells stained with Hoechst, MTG and MitoSOX for detection of superoxide. (J) Superoxide levels of Mdivi-1 treated HeLa cells are elevated (N=6, n≈10800 per condition). (K) Determination of lipid peroxidation by MitoCLox (N=2, n_DMSO_=37, n_Mdivi-1 50 μM 24h_=57, n_Mdivi-1 10 μM 1w_=31). (L) Expression of cardiolipin modifying proteins ALCAT1 and SLP2 in Mdivi-1 treated and control neurons (N=3, n=9). Boxplots indicate median (line), 25th-75th percent percentile (box) and minimum and maximum values (whiskers). Statistics: one-way ANOVA. ***p≤0,001; **p≤0,01; **p≤0,05.

#### Effects of Mdivi-1 on mitochondrial membrane potential

We asked whether a decreased mitochondrial membrane potential ΔΨ_m_ might be the cause of reduced ATP generation. ΔΨ_m_ was determined via fluorescence of tetramethylrhodamine ethyl ester (TMRE), a ΔΨ_m_ *_-_*sensitive dye. We determined ΔΨ_m_ in HeLa cells (Figure S4) and in neurons (Figure 2H). The TMRE signal was normalized on the mitochondrial mass determined by MTG-staining as before (Figure 2G). While short-term (24 h) treatment with Mdivi-1 had no effect on ΔΨ_m_ in neurons, long-term Mdivi-1 treatment (1 w) led to an increased ΔΨ_m_ (Figure 2H). Also, Mdivi-1 treated HeLa cells displayed an increased TMRE signal (Figure S4 A,B). When the uncoupler FCCP was added to the cells a decay in fluorescence intensity was observed. The difference between the ΔΨ_m_ before and after addition of FCCP was calculated. The resulting ΔΔΨ_m_ was also higher in Mdivi-1 conditions. Rotenone treatment on the other hand resulted in a decreased ΔΨ_m_ (Figure S4A).

#### Mdivi-1 treatment results in increased mitochondrial superoxide levels

Impairment of the ETC is often correlated with increased levels of reactive oxygen species (ROS) (Korshunov, Skulachev et al. 1997). We measured superoxide formation via a semi-automated high-throughput imaging approach using the fluorescent dye MitoSOX. HeLa cells with acute Mdivi-1 treatment showed a significant increase in mitochondrial superoxide levels (Figure 2I, J). Following up, the peroxidation state of mitochondrial lipids, in particular cardiolipin, was determined by staining neurons with the fluorescent probe MitoCLox (Lyamzaev, Sumbatyan et al. 2019). Confocal laser scanning imaging revealed no significant differences between Mdivi-1 and control neurons (Figure 2K). Also, the expression of the cardiolipin modifying enzymes Acyl-CoA:lysocardiolipin acyltransferase-1 (ALCAT1) and Stomatin-like protein 2 (SLP2) were not significantly altered due to Mdivi-1 treatment.

### Mdivi-1 reduces complex I assembly and affects supercomplex formation

The reduced respiration rate of mitochondria, the decreased mitochondrial ATP content and the increased superoxide formation indicate a disturbed OXPHOS function of mitochondria in Mdivi-1 treated cells. This suggests a direct effect of Mdivi-1 on OXPHOS complexes. First, we checked Mdivi-1 effects on OXPHOS complex and supercomplex (SC) assembly. Therefore, OXPHOS complexes from isolated mitochondria were separated by Blue Native PAGE (BN-PAGE). Immune-detection of the core subunit NDUFS3 of complex I showed decreased levels of CI-containing SC such as I+III_2_+IV after acute Mdivi-1 treatment (Figure 3A), while the smallest CI-containing SC I+III_2_ showed no significant changes. However, a subcomplex, likely containing an intermediate form of CI, was increased (Figure 3A, B). Immunoblotting of BN-PAGEs for quantification of SC was performed with Anti-NDUFS3 for CI-containing SC (CI^+^-SC, N-respirasome) and subsequent Anti-MTCO1 for SC without CI (CI*^-^ -*SC, Q-respirasome) with the same blot. N- and Q-respirasomes have different molecular weight. To distinguish N- and Q-respirasomes, MTCO1 immune-blotting was first conducted. Next, the membrane was stripped and CI detected by Anti-NDUFS3. This method revealed that the intermediate form likely contains no Complex IV, since no band was detected around 850 kDa when probing the core subunit of Complex IV (MTCO1) as shown in Figure S5C, while the complex was NDUFS3-positive. These findings show that the intermediate SC form is most likely a pre-CI or pre-CI+CIII_2_ in contrast to the pre-CI+CIII_2_+CIV recently proposed (Fang, Ye et al. 2021).

**Figure 3:**
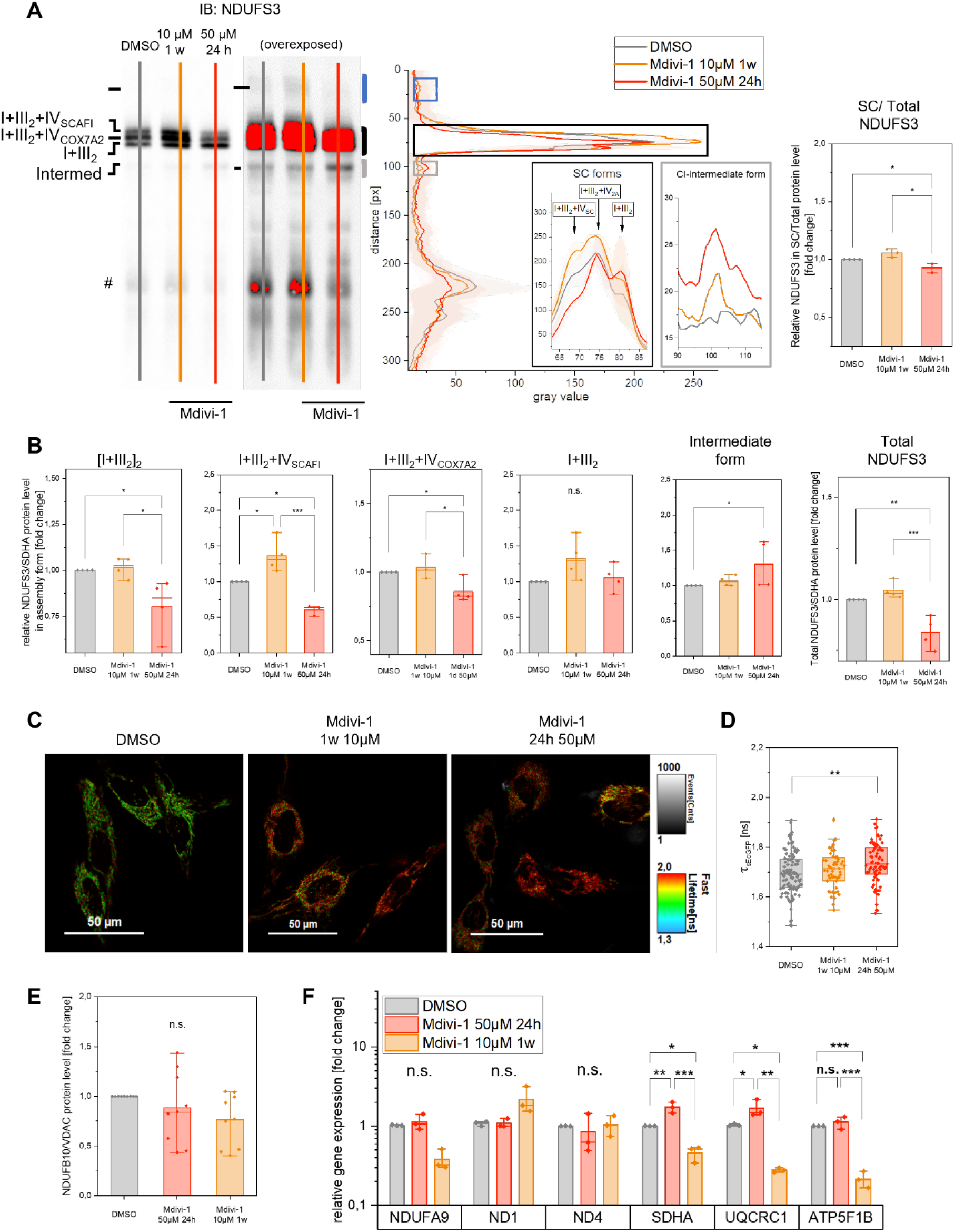
Mdivi-1 treatment impairs CI assembly into supercomplexes. **(A)** Blue-Native separation of proteins and immunoblotting. The line plot of Complex I subunit NDUFS3 shows differences in the assembly of CI and mitochondrial CI-supercomplexes (SC/total) containing CI in between Mdivi-1 treated and control HeLa cells (#=unspecific). (B) Quantification of CI assembly in different isoforms displays alterations between Mdivi-1 and DMSO treated HeLa cells (n=4). I+III_2_ analyzed from blue frame region in (A). (C) Exemplary lifetime images of treated and control HeLa cells expressing the supercomplex sensitive CoxVIIIa-sEcGFP sensor. (D) Lifetime imaging indicates less (CI_1_CIII_2_CIV_1_) supercomplex formation in Mdivi-1 treated HeLa cells (N=3, n_DMSO_=112, n_Mdivi-1,10 µM, 1 w_=55, n_Mdivi-1, 50 µM,24 h_=75 (false color: intensity=gray value, lifetime=rainbow LUT; scale bar: 50 ^µ^m). (E) Protein level of the P-module subunit NDUFB10 is unaltered in Mdivi-1 treated HeLa cells. (N=3, n=9). (F) Gene expression of different OXPHOS subunits is altered due to Mdivi-1 treatment (N=3 independent qPCRs, n=4). Boxplots indicate median (line), 25th-75th percent percentile (box) and minimum and maximum values (whiskers). Statistics: one-way ANOVA. ***p≤0,001; **p≤0,01; **p≤0,05.

Overexposing the immunoblot of the BN-PAGE revealed an assembly containing complex I with higher molecular weight, labeled as [I+III_2_]_2_. However, this could also be the human megacomplex I_2_+III_2_+IV_2_ (Guo, Zong et al. 2017). This assembly form is significantly decreased in acute Mdivi-1 treated HeLa cells compared to DMSO treatment (Figure 3A, blue box in overexposed blot, and Figure 3B). Additionally, an intermediate assembly form (Figure 3A, *intermed*.) was significantly increased in 50 µM Mdivi-1 treated cells, indicating impaired CI assembly into supercomplexes. BN-PAGE with subsequent immunoblotting of the accessory subunit NDUFB10 additionally confirmed this increase of the intermediate form around 850 kDa (Figure S5). The detailed analysis reveals reduced NDUFS3 protein levels in respirasomes I+III_2_+IV_COX7A2_ as well as in I+III_2_+IV_SCAFI_, which includes the supercomplex chaperone SCAFI in acute Mdivi-1 treated cells (Figure 3B). Interestingly, acute treatment decreased this supercomplex form by 40%, while long-term treatment increased the level of I+III_2_+IV_SCAFI_ by about 30%. It is intriguing to speculate that this specific increase in I+III_2_+IV_SCAFI_ is linked to the observed increased ΔΨ_m_ in long-term treated cells. In a complementary assay, CI-containing supercomplex formation was determined by fluorescent lifetime imaging microscopy (FLIM) of a sEcGFP fused to the subunit Cox8a of complex IV (Figure 3C). sEcGFP fused to Cox8a is a supercomplex sensor, changing its fluorescent lifetime depending on its incorporation into supercomplexes (Rieger, Shalaeva et al. 2017). The mean lifetime of sEcGFP in acute Mdivi-1 treated HeLa-Cox8a-sEcGFP cells was 1.73 ± 0.083 ns, while control (DMSO treated) cells displayed a mean sEcGFP-Cox8a lifetime of 1.68 ± 0.085 ns, which was a significant difference (p=0.0018) (Figure 3D). This increase of lifetime in Mdivi-1 treated cells indicates that supercomplex level was decreased, in line with the Blue Native results. The total protein level of NDUFS3 in the BN-PAGE was decreased (Figure 3B), while the NDUFB10 protein level determined by SDS-PAGE (Figure 3E) was not significantly altered. Additionally, mRNA expression levels of the three different Complex I subunits NDUFA9, ND1 and ND4 were not significantly altered after Mdivi-1 treatment (Figure 3F). On the other hand, expression levels of the subunit SDHA of Complex II and UQCRC1 of Complex III were increased in acute Mdivi-1 treated HeLa cells but significantly decreased in long-term Mdivi-1 treated cells compared to the control.

To determine whether the altered complex I assembly affects Q-supercomplex formation (CIII+CIV), the level of supercomplexes containing only complexes IV and III was analyzed in the lower molecular weight region of the BN-PAGE. Indeed, the total amount of the Q-supercomplexes III_2_+IV_2_ and III_2_+IV_1_ was decreased in HeLa cells by acute Mdivi-1 treatment (Figure 4A-B). Also, the mean protein content of complex IV dimers was significantly reduced by 20% (±5% SD), while the amount of monomeric CIV was not altered. This reflects the significance of CI for the stability of CIV and CIV/CIII assembly.

**Figure 4:**
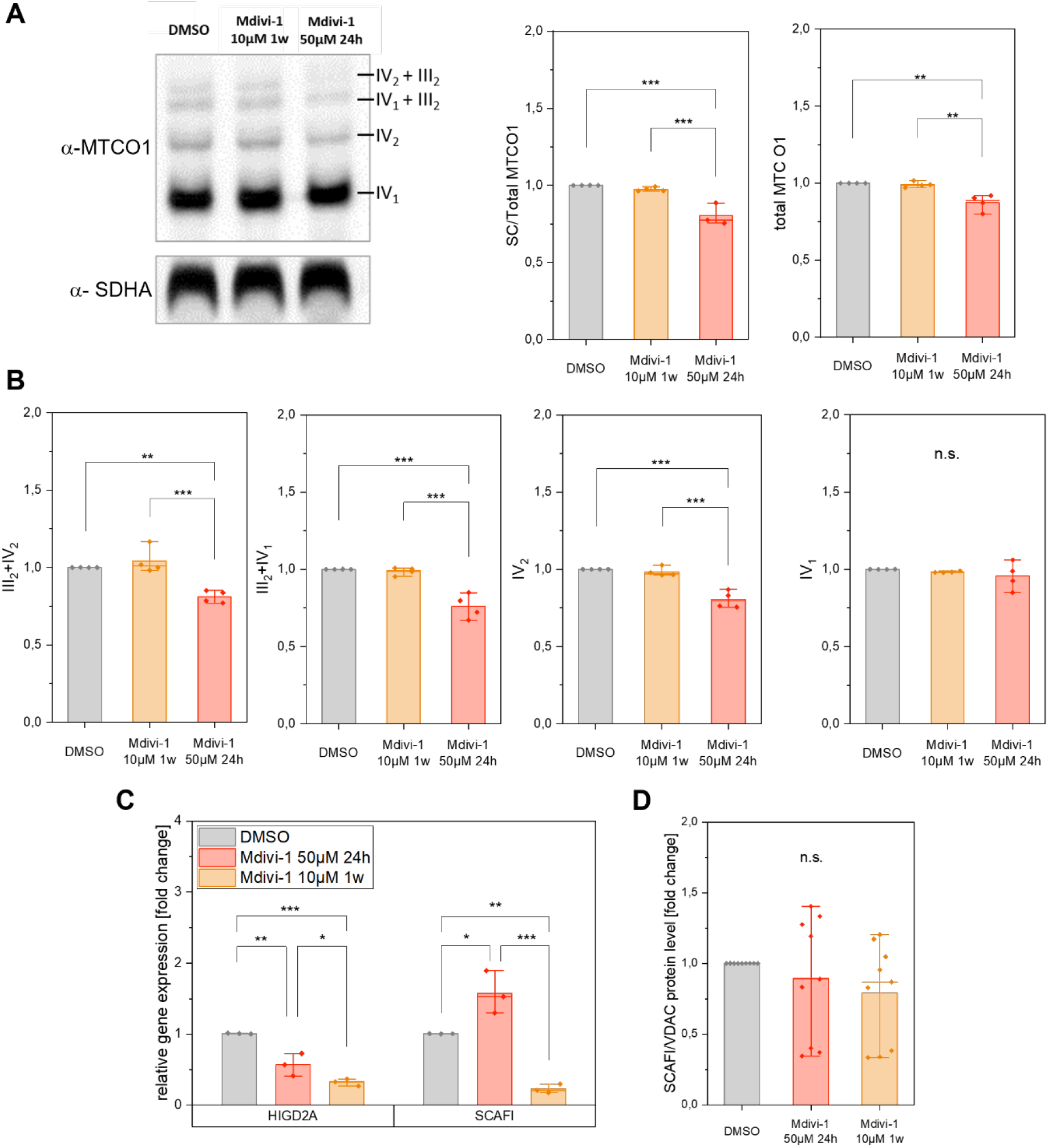
Acute Mdivi-1 treatment decreases the amount of Q-respirasomes. (A) Exemplary immunoblot of BN-PAGE with Mdivi-1 treated HeLa cells. Complex IV probed with α-MTCO1, Complex II probed with α-SDHA). (B) Quantification of supercomplexes reveals decreased supercomplex formation and dimerization of Complex IV (N=4). (C) Expression of supercomplex promoting factors SCAFI and HIGD2A in Mdivi-1 treated cells (N=3, n=12). (D) Protein levels of SCAFI-due to Mdivi-1 treatment (N=3, n=9). Boxplots indicate median (line), 25th-75th percent percentile (box) and minimum and maximum values (whiskers). Statistics: one-way ANOVA with Tukey comparision. ***p≤0,001; **p≤0,01; **p≤0,05.

We checked for changes in Supercomplex Assembly Factor I (SCAFI, also known as COX7A2L), which promotes CIII/CIV interaction (Lapuente-Brun, Moreno-Loshuertos et al. 2013, Vercellino and Sazanov 2021), and Hypoxia Inducible Domain Family Member 2A (HIGD2A). The expression of SCAFI mRNA in HeLa cells treated with Mdivi-1 was significantly increased by the acute treatment and decreased by the long-term treatment. The mRNA expression of HIGD2A was significantly decreased in acute and long-term Mdivi-1 treated cells complexes (Figure 4C). Total SCAFI protein was not significantly changed in Mdivi-1 treated cells, however, this does not report on possible changes of SCAFI bound to respiratory complexes (Figure 4D). Taken together, these data show that acute Mdivi-1 treatment (50 µM, 24 h) impairs the formation of supercomplexes, resulting in the reduction of N- and Q-respirasomes. Long-term treatment (1 w) increases of I+III_2_+IV_SCAFI_ correlated with increased ΔΨ_m._

#### Mdivi-1 inhibits complex I by blocking the quinone binding cavity

We asked whether reduced supercomplex levels in short-term treated Mdivi-1 cells were due to inhibition of complex I by Mdivi as suggested before (Bordt, Clerc et al. 2017). Indeed, it was reported that another inhibitor of complex I, rotenone, resulted in disassembly of supercomplexes (Chapa-Dubocq, Rodriguez-Graciani et al. 2020). To determine the effect of Mdivi-1 on the activity of complex I, respiratory activities were determined as oxygen consumption rates in the presence of substrates and inhibitors. Therefore, cells were permeabilized with digitonin and specific OXPHOS substrates and inhibitors were added. Cells were treated with Mdivi-1 (and DMSO only as control) 24 h prior to the OCR measurement (Figure 5a). Acute Mdivi-1 treatment resulted in a significant decrease in CI+CIII/CIV activity in HeLa cells (Figure 5B) and neurons (Figure S5B). Treatment with 50 ^µ^M Mdivi-1 for 1 h and following washing out resulted in a significant increase of CI+CIII/CIV respiration compared to the treatment with Mdivi-1 persistent in the medium, indicating a regeneration of the complex I respiration and thus reversibility of Mdivi-1 effect. The complex II-dependent respiration (CII+CIII/CIV activity) was not significantly altered upon Mdivi-1 treatment compared to DMSO treated HeLa cells and neurons. Furthermore, a BN-PAGE with isolated mitochondria of Mdivi-1 treated HeLa cells was used for an in-gel activity (IGA) assay. Acutely Mdivi-1 treated HeLa cells displayed a lower intensity of violet bands in the gel indicating reduced CI activity (Figure 5C).

**Figure 5:**
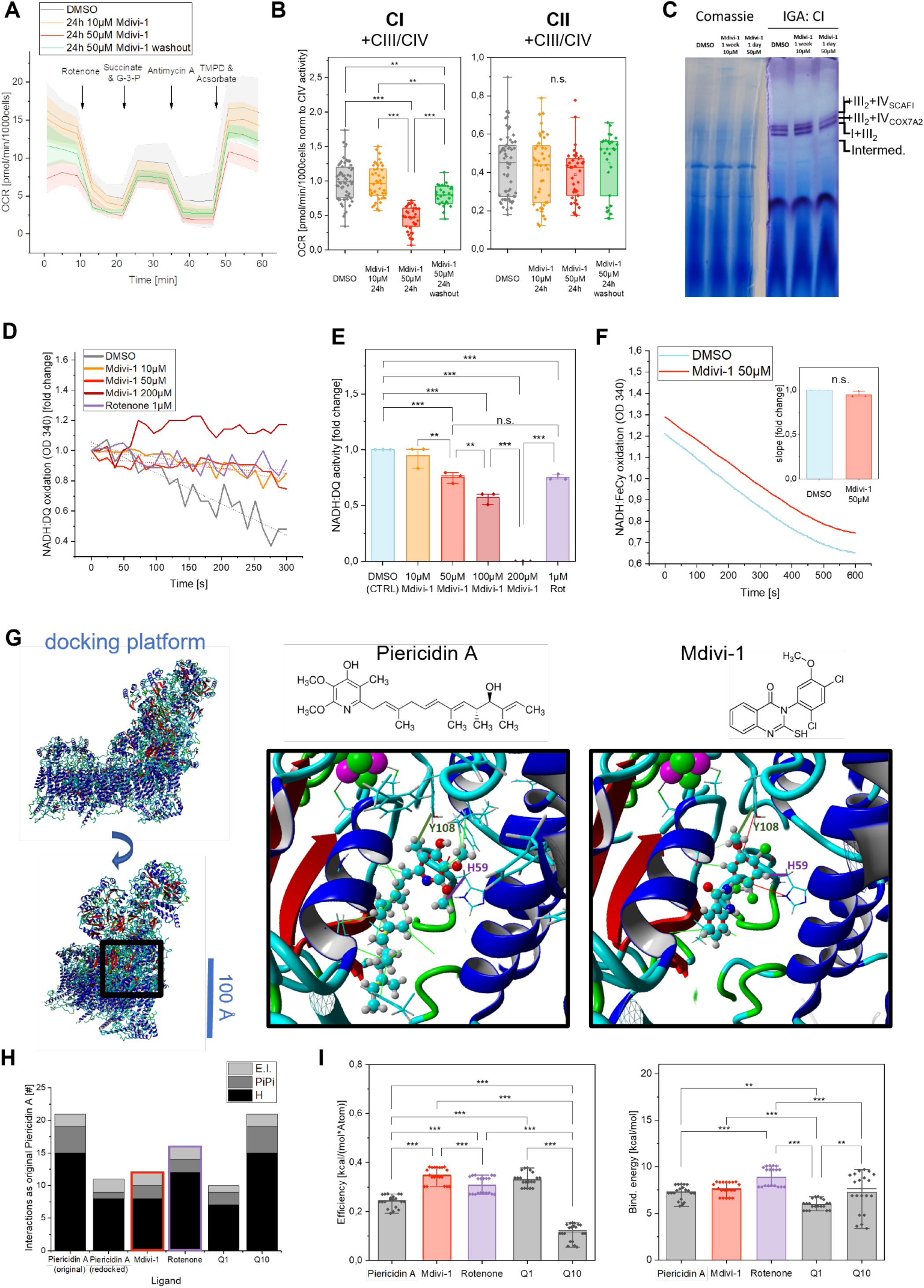
Mdivi-1 acts as a local inhibitor in the Q-cavity of Complex I. (A-B) Determination of respiratory complex activities in HeLa cells by oxygen consumption rates (N=3, n_DMSO_=41, n_Mdivi-1,10 µM, 1 w_ =41, n_Mdivi-1, 50 µM,24 h_ =35, n_Mdivi-1 50_ ^µ^_M 24h washout after 1h_=27). (C) BN-PAGE gel shows reduced complex I in-gel activity in isolated mitochondria from acute Mdivi-1 treated HeLa cells. (D) Complex I activity determined by NADH:DQ oxidoreduction activity after treatment with Mdivi-1 and Rotenone in isolated mitochondria of HeLa cells (E) Complex I activity determined by NADH:DQ oxidoreduction activity after treatment with Mdivi-1 or Rotenone in isolated mitochondria of NPC (N=3). (F) Effect of Mdivi-1 treatment on complex I NADH:FeCy oxidoreduction activity in isolated NPC mitochondria determined by (N=3). (G) Representative docking poses with residue interactions of piericidin A and Mdivi-1 in complex I [*Mus musculus*] (6ZTQ) generated with Autodock (Hydrophobic interactions=light green, PiPi interaction=red, interaction with Tyrosine 108=dark green, interaction with Histidin 59=violet). (H) Number of interactions in best docking poses of Autodock experiment identical to the piericidin control in the inhibitor-bound cryo-EM structure. (I) Efficiency and binding energy of all dockings with different ligands (N=4 docking processes, n=25 docking poses). Boxplots indicate median (line), 25th-75th percent percentile (box) and minimum and maximum values (whiskers). Statistics: one-way ANOVA. ***p≤0,001; **p≤0,01; **p≤0,05.

To test whether Mdivi-1 is a direct inhibitor of complex I, mitochondria of HeLa wildtype cells were isolated and the mitochondrial membrane was disrupted to dissociate individual mitochondrial complexes. The enzyme activity of total complex I was determined by time-dependent NADH oxidation (Figure 5D). NADH was used as electron donor and decyl-ubiquinone (DQ) was used as electron acceptor in the activity buffer, while antimycin A was used to block further electron transfer to Complex III. NADH:DQ activity in mitochondria of HeLa (Figure 5D) and NPCs (Figure 5E) were measured in NADH:DQ activity buffer containing different concentrations of Mdivi-1 and rotenone. Mdivi-1 concentrations of at least 50 ^µ^M showed a significant decrease in complex I activity compared to control. This led to the conclusion that Mdivi-1 is a direct inhibitor of complex I. The NADH:DQ activity of 1 ^µ^M rotenone and 50 ^µ^M Mdivi-1 did not differ significantly, indicating a similar inhibitory effect. To determine whether Mdivi-1 interacts in the Q-cavity or with the peripheral arm of complex I, the NADH oxidation activity of the enzyme activity was determined. For that, ferricyanide was used as an acceptor for electrons. The NADH:FeCy oxidation of NPC mitochondria was not significantly altered by addition of 50 ^µ^M Mdivi-1 to the activity buffer (Figure 5F). This suggests that blocking the quinone binding cavity of complex I was responsible for the inhibition of CI.

#### The ubiquinone reduction activity of complex I is inhibited by Mdivi-1

To address, whether the Q-cavity binds Mdivi-1, an *in-silico* approach was used to determine interactions and binding energies of Mdivi-1, ubiquinones and known inhibitors of complex I. The cryo-EM structure of piercidin-binding complex I from *Mus musculus* (6ZTQ) was used as a docking platform. First, the molecular structure of the inhibitor piercidin A was manually removed from the overall structure (6ZTQ). Different molecular docking algorithms were used for a global docking with the ligands piericidin A, Mdivi-1, rotenone, ubiquinone (Q1) and ubiquinone-10 (Q10). The best docking poses of the re-introduced piericidin A (CTRL) and Mdivi-1 in the cropped version of 6ZTQ generated by Autodock are shown in Figure 5G. The demonstrated interactions with residues in the Q-cavity were analyzed by the number of identical interactions of piericidin A found in the original cryo-EM structure of complex I co-crystallized with piericidin A. The original piericidin A contained 21 interactions, while the docking pose generated by redocking piericidin A (CTRL) contained 11 interactions. The docking of all ligands, including Q10 but not Q1, involved two essential interactions (Bridges, Fedor et al. 2020) (Murai 2020) that are critical for binding to this complex I cavity (Figure 5H). The total number of identical interactions was higher for the ligands Mdivi-1, rotenone and Q10 compared to piericidin A. In an analogous docking experiment using the complex I cryo-EM structure of *Bos taurus* (5LDW), first a structural alignment was conducted to determine the homology between the two structures based on their shapes and three-dimensional conformations. Structure alignment of the full complex I structures of 6ZTQ and 5LDW showed a root-mean-square deviation (RMSD) of atomic positions of 4.32 Å (Figure S6A), which is low for a large protein complex like complex I. Docking of inhibitors to complex I from *Bos taurus* (5LDW) showed fewer interactions (Figure S6B) and lower binding energies (Figure S6C) for each ligand. The overall efficiencies and binding energies of docking calculations using Autodock algorithms and all structures of each host organism were pooled for final comparison. The mean efficiency for all docking poses determined in YASARA was significantly increased for the ligands Mdivi-1 and rotenone compared to piericidin A control (Figure 5I) whereby Mdivi-1 showed higher efficiency than rotenone. On the other hand, Mdivi-1 poses showed the same binding energy as rotenone (Figure 5I). Together with the *in vitro* findings these experiments indicate that Mdivi-1 acts as a local inhibitor in the Q-cavity of complex I similar to rotenone.

### Long-term treatment with Mdivi-1 significantly reduces synaptic activity in neurons

To determine the significance of Mdivi-1 inhibition of complex I for neuronal activity, we conducted electrophysiological measurements of stimulated neurons via Microelectrode Arrays (MEA). Because of the mainly glutamatergic neuronal cell culture, pharmacological stimulation was performed with glutamate/glycine (each 100 µM) (Figure 6A). The number of peaks per minute was significantly decreased by 38% in long-term Mdivi-1 treated neurons compared to control neurons (Figure 6B). Acute Mdivi-1 treatment of neurons led to a non-significant decrease by 11%. Figure 6C plots the mean peak amplitude in time intervals of 5 s. Neurons with long-term Mdivi-1 treatment had a reduced mean amplitude of 0.16 mV (± 0.05 SD) compared to the DMSO mean amplitude of 0.43 mV (± 0.05 SD). Both conditions showed a relatively constant amplitude, while the amplitude of acute Mdivi-1 treated neurons conveys a decreasing trend (Figure 6C). To test, whether vesicle fusion was involved in deterioration, we determined two marker proteins of the pre-synapse: synaptophysin (SYP) and syntaxin 4 (STX4). SYP is also known as the major synaptic vesicle protein p38. STX4 is part of the SNARE complex, which induces the fusion of synaptic vesicles with presynaptic terminals. Immunoblotting of SYP revealed a significant increase of the protein in differentiated neurons compared to non-differentiated NPC. Cells with acute Mdivi-1 treatment showed a tendency towards an elevated SYP level compared to DMSO treated neurons (Figure 6D), indicating that Mdivi-1 did not affect the protein levels of SYP. However, the expression of STX4 was significantly reduced in long-term Mdivi-1 treated neurons (Figure 6E). This could cause an impaired fusion of synaptic vesicles with the membrane after stimuli. Since vesicle fusion is Ca^2+^-dependent, we qualitatively monitored Ca^2+^-dynamics in NPC-derived neurons stained with the calcium-indicator dye Fura-2. Pharmacological activation clearly evoked a response visible as calcium transients, demonstrating that neuronal activity was monitored (Figure S6A). Calcium traces showed no significant difference of basal activity due to Mdivi-1 treatment, but the stimulated cells exhibited significantly less calcium uptake in neurons, that were treated with 10 µM Mdivi-1 for one week during the differentiation process (Figure S6B).

**Figure 6:**
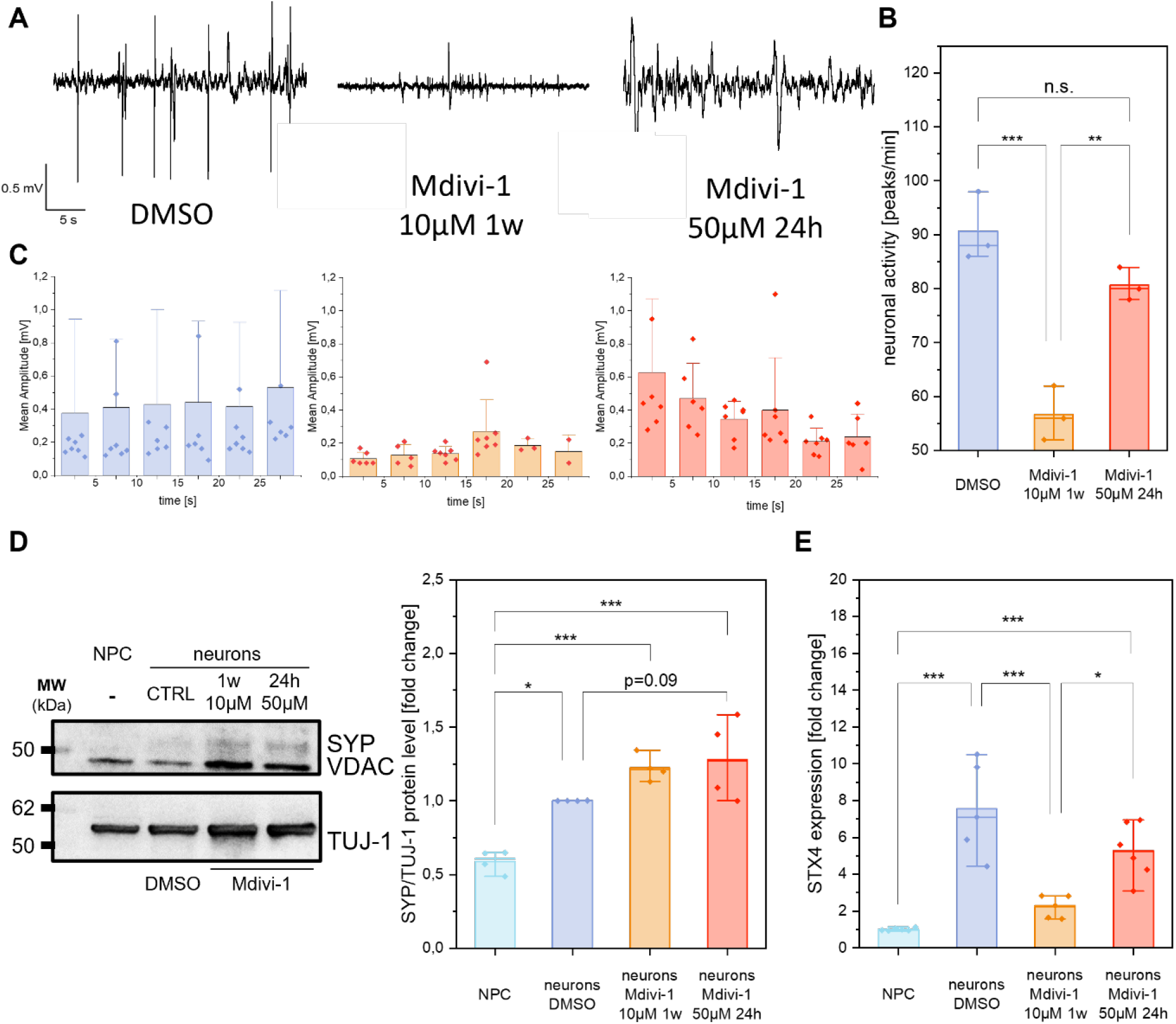
Mdivi-1 reduces neuronal function. (A) Exemplary MEA traces after stimulation with glutamate/glycine (100 ^µ^M each). (B,C) Peak number and amplitude in control and Mdivi-1 treated neurons (N=2 differentiations, n=9, ANOVA). (D) Quantification of synaptophysin (SYP) protein levels in control and Mdivi-1 treated NPC-derived neurons (N=2, n=4, ANOVA). (E) Gene expression of Syntaxin 4 is decreased due to Mdivi-1 treatment (N=2, n=6, ANOVA). Boxplots indicate median (line), 25th-75th percent percentile (box) and minimum and maximum values (whiskers). Statistics: one-way ANOVA correction. ***p≤0,001; **p≤0,01; **p≤0,05.

### Mdivi-1 alters the cellular and mitochondrial calcium homeostasis

To investigate Mdivi-1 effects on Ca2+ homeostasis, fluorescent FRET-based biosensors of the Chameleon family were used. The tests were exemplary done in HeLa cells. The basal cytosolic calcium level was not altered in Mdivi-1 treated cells compared to DMSO control. Cytosolic calcium uptake was studied after stimulation with extracellular calcium chloride (2 mM) (Figure 7A). To study the Mdivi-1 effect, HeLa cells were treated with 10 ^µ^M and 50 ^µ^M Mdivi-1 for 24h. The mean cytosolic calcium uptake of 10 ^µ^M pretreated cells was decreased by 55%, while it was decreased by 75% in 50 ^µ^M pretreated cells (Figure 7B). To determine effects of Mdivi-1 on the mitochondrial calcium levels, cells were transfected with a calcium sensor fused to a targeting sequence for the mitochondrial matrix (Figure 7C). The mitochondrial calcium level of Mdivi-1- and rotenone-treated HeLa cells were significantly elevated (Figure 7D).

**Figure 7:**
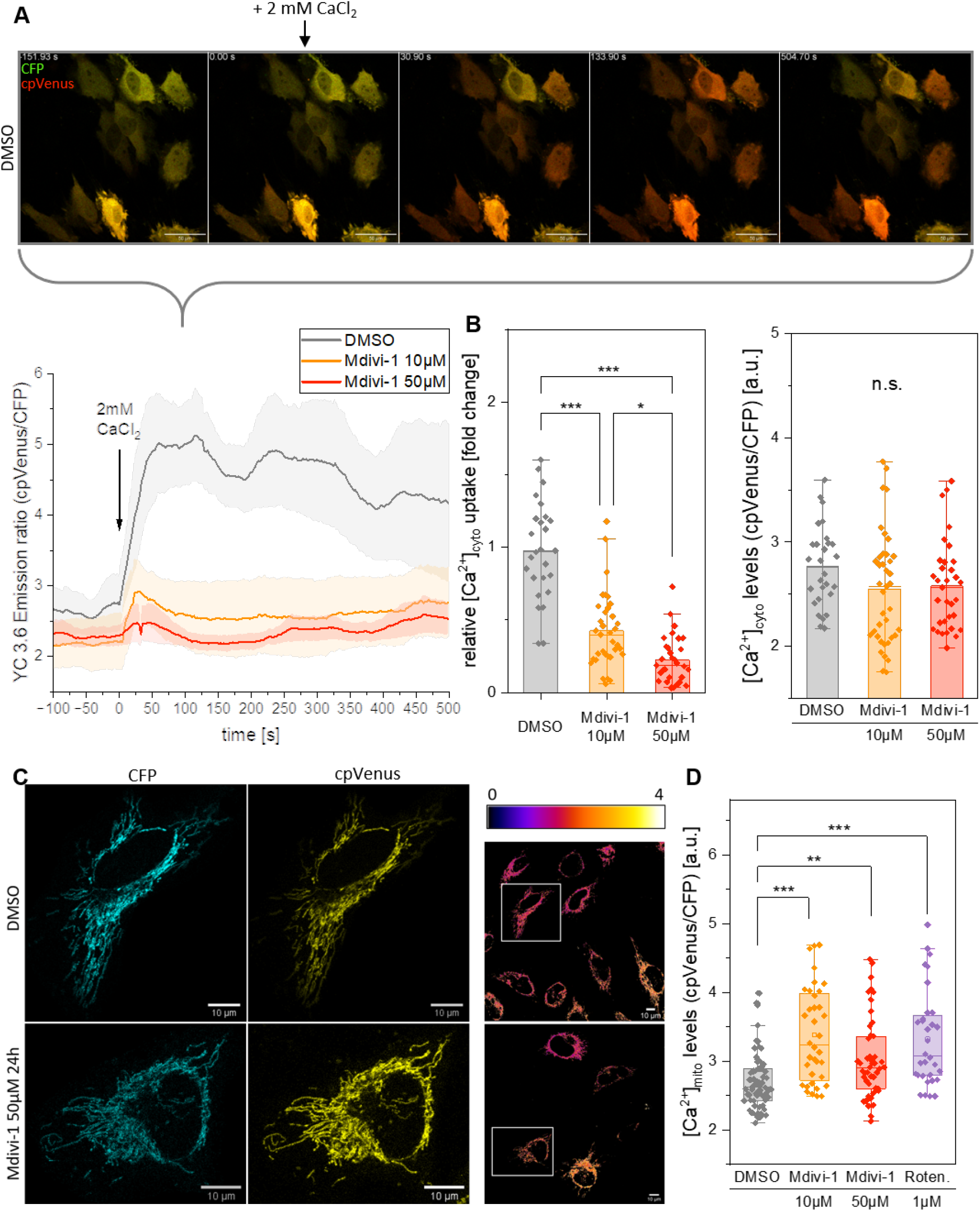
Mdivi-1 alters cellular and mitochondrial calcium homeostasis. (A) Representative time series of a [Ca^2+^]_cyto_ uptake experiment monitored via the YC 3.6 biosensor in transiently transfected HeLa (false color: green=CFP, red=cpVenus) and ratio-metric quantification of [Ca^2+^]_cyto_ uptake experiment with one biological replicate (scale bar: 50 ^µ^m). (B) Basal cytosolic calcium uptake is impaired in Mdivi-1-treated HeLa cells (N=4, n_DMSO_=26, n_Mdivi-1,10 µM, 24 h_ =35, n_Mdivi-1, 50 µM,24 h_ =31, KW). (C) Exemplary images of mt4D3-cpV transfected HeLa cells (false color: cyan=CFP, yellow=cpVenus; ratiometric image with fire LUT; scale bar: 10 ^µ^m). (D) Mdivi-1 treatment leads to elevated mitochondrial calcium levels (N=3, n_DMSO_=67, n_Mdivi-1,10 µM, 24 h_ =36, n_Mdivi-1, 50 µM,24 h_ =47, n_Rotenone_=32, KW). Boxplots indicate median (line), 25th-75th percent percentile (box) and minimum and maximum values (whiskers). Statistics: one-way ANOVA. ***p≤0,001; **p≤0,01; **p≤0,05.

## Discussion

Mdivi-1 has been discussed as a potential therapeutic treatment for neurodegenerative diseases (Liu, Song et al. 2022). However, the mechanism of action is still not fully understood and recent data suggest an inhibitory effect not only on Drp1, the mitochondrial fission factor, but on mitochondrial bioenergetics (Bordt, Clerc et al. 2017, Ruiz, Alberdi et al. 2018, Bordt, Zhang et al. 2022). It. We here showed in detail that Mdivi-1 is a Complex I inhibitor that binds to the I_Q_-site similar as rotenone. Mdivi-1 treatment resulted in reduced CI activity and oxygen consumption rates, decreased mitochondrial ATP levels (Figure 5A-E) and impaired neuronal activity (Figure 6), involving altered calcium dynamics (Figure 7, Figure S7).

As a mechanism of action, we showed that Mdivi-1 specifically inhibits the ubiquinone reduction in respiratory Complex I (I_Q_-site), while NADH:FeCy oxidation was not affected. NADH:DQ oxidations rates were reduced in Mdivi1-treated mitochondria. The Q-cavity is a target of a variety of structurally different CI inhibitors, in particular of rotenoids (e.g. rotenone) (Degli Esposti 1998). The potency of rotenoids to bind to the I_Q_-side is due to a specific spatial organization of hydrogen-bond acceptable methoxy oxygens that allow the tight fitting into the binding site and provide a bending axis between thermodynamically stable phenol rings. The structure of Mdivi-1, chemically 3-(2,4-dchloro-5-methoxyphenyl)-2-sulfanyl-4(3H)-quinazolinone, contains three aromatic rings which are also not in one plane. The bond between the quinazoline moiety and the phenyl provides the bending axis that is required for the stable positioning of Mdivi-1 in the I_Q_-side of Complex I (Figure 5G). The oxygen atom of the hydroxyl group and the chloride ion of the first aromatic ring in Mdivi-1 provide similar binding possibilities as the two methoxyl groups on the A-ring of rotenone. The computational interaction analysis revealed additional hydrogen bonds Met69 (NDUFS7), Met70 (NDUFS7), Phe86 (NDUFS7) and Thr156 (NDUFS2) that are reported to further stabilize the inhibitors in the binding site (Bridges, Fedor et al. 2020).

Physiological measurements confirmed that Mdivi-1 is a Complex I inhibitor that inhibits oxidative metabolism and increases ROS. Similar effects on bioenergetics were observed in DRP1-deficient H460 cells before (Dai, Wang et al. 2020). This shows that Mdivi-1 affects bioenergetic function independent of mitochondrial fission factor Drp1. Since 50 µM Mdivi-1 had comparable effects to 1 µM rotenone, one of the strongest inhibitors of the Complex I I_Q_-binding site (Figure 5H), a higher *K*_I_ of Mdivi-1 can be assumed. Mdivi-1 exclusively inhibited CI, since CII-dependent respiration was not altered. We found increased ΔΨ_m_ and decreased ATP levels in Mdivi-1 treated cells. The increase in ΔΨ_m_ could be due to reverse ATP synthase activity, induced by the compromised ETC. This would also explain lower mitochondrial ATP levels. In addition, ROS levels increased (Figure 2I, J). These effects are similar as it was earlier observed for Rotenone (Forkink, Manjeri et al. 2014). With respect to the mechanism of inhibition, Mdivi-1 and Rotenone thus are similar, both increase ROS levels. This is typical for A-class type inhibitors of the I_Q_-site of Complex I (Fato, Bergamini et al. 2009, Rieger, Thierbach et al. 2020). In contrast, Class B inhibitors (e.g. stigmatellin) do not affect ROS production, since they block the electron transfer from the iron-sulfur cluster N2 to ubiquinone in a different way. As no change in Complex I activity was observed at the FMN site (Figure 5F), Mdivi-1 is not an I_F_ inhibitor. It was suggested that binding of Mdivi-1 to Complex I is reversible (Bordt, Clerc et al. 2017), but we found only partial regeneration of Complex I dependent respiration after removal of Mdivi-1. The incomplete regeneration could be due to Complex I degradation and supercomplex (SC) disassembly, respectively, as discussed in the following paragraph.

### Potential mechanisms for the discovered alterations in SC formation

Acute Mdivi-1 treatment decreased the formation of respiratory SCs containing CI, the N-respirasomes (Figure 3), but also SCs without CI, the Q-respirasomes (Figure 4). The relative mRNA expression of SC assembly factor HIGD2A was significantly decreased by Mdivi-1 treatment. HIGD2A promotes either isolated CIV assembly through addition of the Cox3 module or associates with CI+CIII_2_ supercomplexes to add CIV for the formation of a respirasome (Timon-Gomez, Garlich et al. 2020). Reduced interaction between SC I+III_2_ and CIV and CVI_2_, respectively, can explain the increase of I+III_2_ compared to respirasomes as shown in the SC line plot of Figure 3A. Transcriptional regulation of HIGD2A function is a regulator of respiratory supercomplexes assembly in response to hypoxia, cellular metabolism and cell cycle: knock out of HIGD2A in the murine skeleton muscle C2C12 cell line resulted in less SC, higher OPA1 levels, increased [ROS]_mito_ and increased ΔΨ_m_ (Salazar, Elorza et al. 2019). These effects match our findings. It is suggested, that SCAFI stabilizes SC without CI (Perez-Perez, Lobo-Jarne et al. 2016). The observed protein level of the assembly factor SCAFI was not significantly decreased, however the standard deviation in between replicates of the immunoblotting of SCAFI was very high and the total protein level does not represent the amount of SCAFI protein in the assembled complexes. In contrast to the decreased SC without CI, the mRNA expression of SCAFI was increased in HeLa cells, which might be a compensation due to the decrease of all supercomplexes. A SC form denoted with [I+III_2_]_2_ of higher molecular weight was observed at ≈ 1300 kDa showing either a respirasome with a trimer of CIV (I+III_2_+IV_3_), or I_2_+III_2_ (Jha, Wang et al. 2016) or the human mitochondrial megacomplexes (Guo, Zong et al. 2017). To further dissect the exact composition of complexes in the quantified band, quantitative mass spec analysis would be required (Gonzalez-Franquesa, Stocks et al. 2021).

The separation of mitochondrial complexes by BN-PAGE and subsequent immunoblotting of complex I subunits NDUFS3 and NDUFB10 revealed an increase of an intermediate form with a molecular weight of approximately 850 kDa (Figure 3 and Figure 4). Monomeric complex I is hardly found in human mitochondria. Therefore the intermediate form can be either the smallest CI containing SC without the last assembly factors (pre-I+III_2_) (Moreno-Lastres, Fontanesi et al. 2012), a pre-CI, which is a subassembly with a molecular weight of ≈ 830 kDa (Vartak, Semwal et al. 2014) or a pre-respirasome subcomplex, containing not fully assembled CI. The following sections will discuss potential reasons for the increase of this intermediate form and will also discuss the different pathways for complex I assembly.

#### CI perturbation reduces the level of N-respirasomes

Recent studies have shown that knockout of NDUFB10, which is an accessory subunit of CI stabilizing the P-arm of the complex, results in incomplete assembly of Complex I (Stroud, Surgenor et al. 2016, Arroum, Borowski et al. 2023). The loss of subunits of the N- and P-module ultimately led to the loss of Complex I and respiratory supercomplexes. Assembly analysis of the CI-containing supercomplexes in a study with different knockout cell lines of CI accessory subunits (e.g. NDUFA8-KO, NDUFS5-KO, NDUFC1-KO) revealed the loss of supercomplexes by BN-PAGE (Stroud et al. 2016). Those studies provide evidence that fully assembled CI is required for SC formation, as previously described (Moreno-Lastres et al. 2012). Furthermore, it is suggested that CI assembles in the absence of CIII but is unstable, and inhibition of CIII activity does not affect CI assembly (Acín-Pérez, Bayona-Bafaluy et al. 2004). Since an intermediate form of CI (similar to pre-CI proposed in (Vartak, Semwal et al. 2014)) was found in Mdivi-1 treated cells, this led us conclude that impairment in CI assembly results in a decrease of supercomplexes.

#### The decrease of supercomplexes (with and without CI) leads to CI destabilization

The NDUFB10-KO as well as a ND6-KO cell line of CI completely prevented the formation of N-respirasomes, but still allowed the-formation of the supercomplex III+IV (Acin-Perez, Fernandez-Silva et al. 2008). Here, we found also a decrease of supercomplexes without CI (Q-respirasome). The cooperative model suggests that CI builds up as a pre-CI of ≈ 830 kDa, with the binding site of the N-module subunit being occupied by NDUFA12 to stabilize the pre-CI. In addition, CIV associates with CIII, which in turn binds to the pre-CI (Guan, Zhao et al. 2022). Finally, NDUFA12 is exchanged to assemble the N-module to provide a functional respirasome. According to this model, the impaired connection of CIII with CIV can hinder the biogenesis of CI. This argumentation is also relevant for another proposed assembly pathway, which describes a similar cooperative model, but an earlier association of CIII with a membrane arm of CI (Fang, Ye et al. 2021). A decrease of N-respirasomes would decrease the integrity and stability of CI. The N-module is assembled as the last step in these two models, but the observed intermediate form shows slight NADH-oxidation by CI in-gel activity (Figure 5C), which suggests that the destabilized CI form still contains the N-module. The assembly of CI into supercomplexes regulates ROS production and may contribute to the bioenergetic differences between neurons and astrocytes (Lopez-Fabuel, Le Douce et al. 2016). A decrease of all SC forms increases the ROS production, which is what we observed.

CI inhibition leads to a conformational change in the protein, altering the closed state and the angle of the membrane arm to the peripheral arm, and additionally preventing its function. Persistent binding of Mdivi-1 to Complex I could therefore alter the interaction with Complexes III and IV and thus contribute to N-respirasome disassembly and CI instability. Whether CI inhibition directly affects CI assembly or SC formation has not been reported in the literature and was not further investigated in this study.

### Long-term Mdivi-1 inhibition of CI and disturbance of ETC function impairs neuronal activity

Long-term Mdivi-1 treatment during differentiation resulted in reduced electrical activity of neurons, as exhibited in fewer electric spikes per minute using a MEA assay (Figure 6A). Mitochondria control neuronal activity mainly by providing ATP and mediating calcium signaling required for vesicular exocytosis, endocytosis and vesicle recycling, as well as for powering synaptic transmission. The impaired ETC activity due to decreased Complex I activity and reduced N- and Q-respirasome levels resulted in decreased ATP production, although the ΔΨ_m_ was increased. This strongly indicates a lower P/O-Quotient (coupling of respiration and ATP synthesis) in Mdivi-1 treated cells and thus reduced efficiency of ATP production, which eventually led to energy starvation of the neurons. We further found that long-term treatment with Mdivi-1 (10 µM, 1 week) resulted in an attenuated level of cytosolic calcium in HeLa cells and neurons that were stimulated with glycine/glutamate (Figure 7B and Supplementary Figure S7). This can be related to decreased ATP, be due to an impaired function (or decreased protein level) of neuronal Voltage-gated calcium channels (VGCCs) or be due to an altered intra-cellular calcium buffering. Indeed, we found that [Ca^2+^]_mito_ levels were increased in HeLa cells. We assume that mitochondrial calcium buffering is also increased in neurons and thus the calcium homeostasis and dynamics of neurons is altered due Mdivi-1 treatment, which is in line with a previous report (Ruiz, Alberdi et al. 2018). This study showed that short Mdivi-1 treatment (50 µM, 1h) induced a reduction of cellular and mitochondrial Ca^2+^ uptake, when cells were exposed to NMDA or AMPA/CTZ.

Another reason for the reduced activity of long-term Mdivi-1-treated neurons may be the impairment of synapse formation at a later stage of differentiation. We found no change of the total presynaptic vesicle-related SYP protein level after short- or long-term Mdivi-1 treatment, though (Figure 6D). However, we found a reduced Syntaxin 4 expression (Figure 6E). Syntaxins bind synaptotagmin in a calcium-dependent fashion and interact with voltage dependent calcium and potassium channels. Direct syntaxin-channel interaction links the vesicle fusion machinery and the gates of [Ca^2+^]_cyto_ entry during depolarization of the presynaptic axonal boutons (Han, Wang et al. 2004). Our current hypothesis is that long-term Mdivi-1 treatment leads to reduced neuronal function due to impaired calcium-dependent vesicle fusion, which reduces exocytosis.

## Conclusion

In view of the results presented here, a possible therapeutic application of Mdivi-1 must take into account the dose- and time-dependent effects on mitochondrial energy metabolism. Complex I of the respiratory chain is central to OXPHOS activity, ATP formation and calcium homeostasis and its inhibition has effects on neuronal activity

## Material and Methods

## Material

### Primers, Antibodies, Kits, Chemicals

**Table 1:**
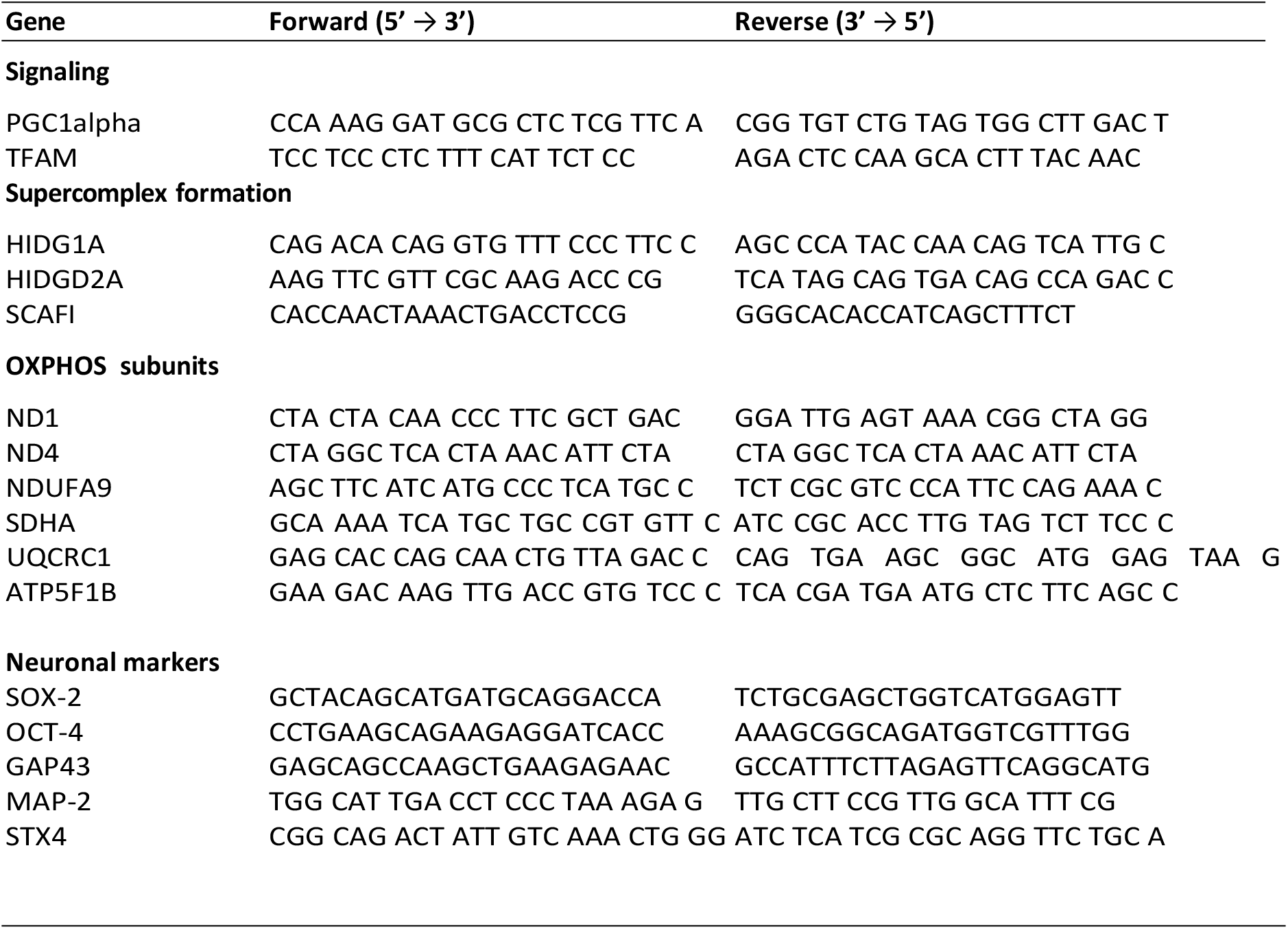
List of used primers for quantitative PCR (qPCR).

**Table 2:** List of used primary antibodies for Immunofluorscence (IF) and Immunoblotting (IB).

| Primary Antibody | Manufacturer | source | Dilution (IF) | Dilution (IB) |
| --- | --- | --- | --- | --- |
| Anti-ATP5i | abcam (ab12241) | rb | 1:200 | - |
| Anti-CHAT | Proteintech (20747-1-AP) | rb | 1:200 | - |
| Anti-MAP2 | Invitrogen (PA5-17646) | rb | 1:400 | - |
| Anti-MTCO1 | Invitrogen (459600) | ms | - | 1:2000 |
| Anti-NDUFS3 | Abcam (ab183733) | rb | - | 1:2000 |
| Anti-NDUFB10 | Abcam (ab196019) | rb | - | 1:5000 |
| Anti-COX7A2L [SCAFI] | St Johns Laboratory (STJ110597) | rb | - | 1:1000 |
| Anti-SDHA | Abcam (Ab14715) | ms | - | 1:2000 |
| Anti-SYP | Santa Cruz Biotechnology (sc-17750) | ms | - | 1:2000 |
| Anti-TH (H-196) | Santa Cruz Biotechnology (sc-14007) | rb | 1:200 | - |
| Anti-GHITM [TMBIM5] | Proteintech (16296-1-AP) | rb | - | 1:1000 |
| Anti-TUJ-1 | GRP (GT11710) | ms | 1:400 | 1:2000 |
| Anti-VDAC 1/2 | Proteintech (10866-1-AP) | rb | - | 1:2000 |
| Anti-Acetylated Tubulin | Sigma (T7451) | ms | 1:4000 | - |

**Table 3:**
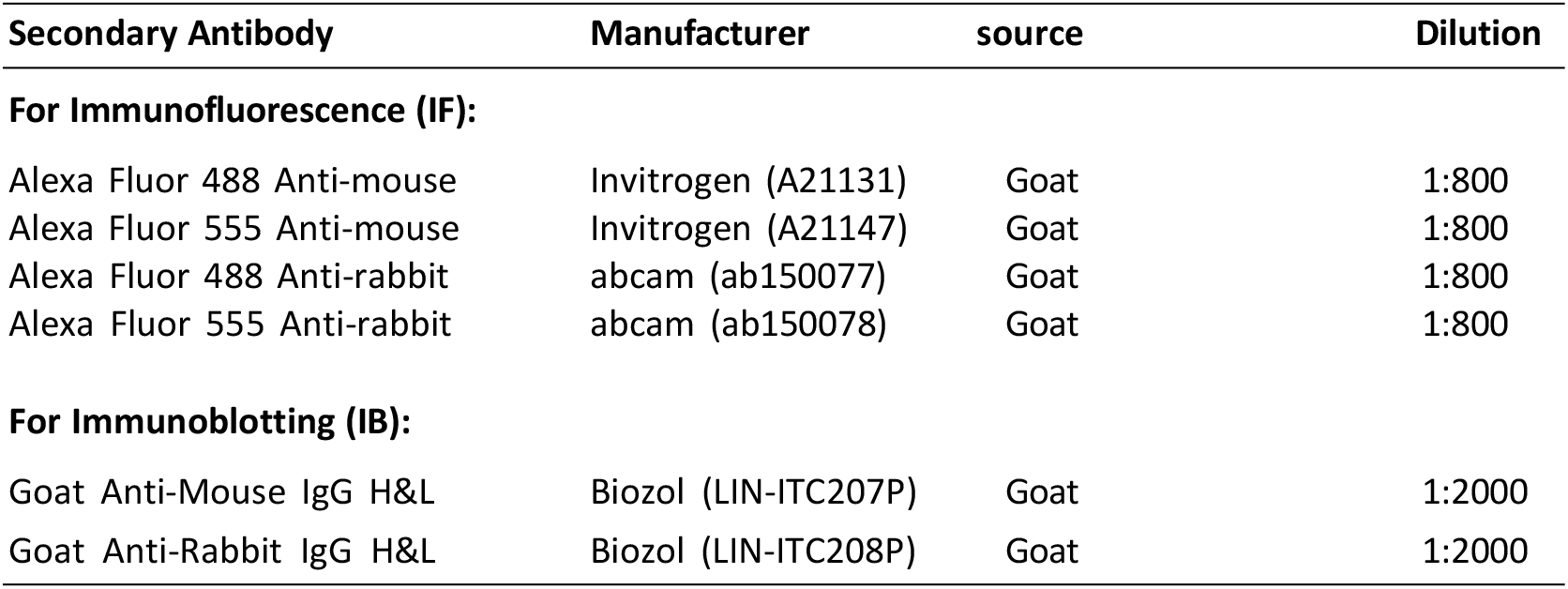
List of used secondary antibodies.

**Table 4:** List of dyes used.

| Dye | Manufacturer | (Cat. No) |
| --- | --- | --- |
| BioTracker ATP-Red Live cell dye | Sigma-Aldrich | (SCT045) |
| Fura-2 AM | Thermo Fisher | (F1221) |
| FxCycle PI/RNase Solution | Thermo Scientific | (F10797) |
| Janelia Fluor HaloTag Ligand 646 (JF HTL 646) | Promega | (GA1120) |
| Mito Tracker Deep Red FM (MTDR) | Thermo Fisher | (M22426) |
| Mito Tracker Green FM (MTG) | Thermo Fisher | (M7514) |
| Mitochondrial Superoxide Indicator (MitoSox) | Thermo Fisher | (M36008) |
| MitoClox | gift from Boris Chernyak |  |
| Tetramethylrhodamin-methylester (TMRE) | Thermo Fisher | (T669) |

**Table 5:** List of used kits.

| Name | Manufacturer |
| --- | --- |
| Monarch Total RNA Miniprep Kit | New England BioLabs (T2010S) |
| Seahorse XF Cell Mito Stress Test Kit | Agilent (103010-100) |
| RevertAid First Strand cDNA Synthesis Kit | Thermo Scientific (K1621) |
| SuperSignal West Pico PLUS Chemilum. Substrate | Thermo Scientific (S8761) |
| TurboFectin 8.0 | OriGene (TF81001) |

## Methods

### Cell lines

#### HeLa cells

Hela cells were purchased from the Leibniz Institute DSMZ-German Collection of Microorganisms and

Cell Cultures Hela cells were cultured in T25 flasks and maintained at 5% CO_2_ and 37°C in supplemented MEM-medium (Minimum essential medium Eagle, Sigma-Aldrich, M2279; 10 % FBS supreme, PAN BioTech P30-3031; 1% HEPES, Sigma-Aldrich H0887-100ML, Ala-Gln, Sigma-Aldrich G8541-100ML, 1%; MEM non-essential amino acid solution, Sigma-Aldrich, M7145-100ML, 1%). Cells were split every 2-3 days and were not allowed to exceed 90% confluency.

#### Neuronal progenitor cells

Neuronal progenitor cells (NPC) were a kind gift of Prof. Thomas Gasser, Neurologische Universitätsklinik Tübingen, Germany.Neuronal progenitor cells (NPC) were cultured on 1% [(v/v) in KO-DMEM/F-12, Thermo Fisher] matrigel (BD Biosciences)-coated 6-well plates (Sarstedt) in Neuronal Keeping Medium (NKM), composed of N2B27 medium with addition of the small molecules smoothened agonist (SAG, 0.5 ^µ^M, Cayman Chemical) and CHIR 99021 (3 ^µ^M, Axon MedChem) and 150 ^µ^M ascorbic acid (Sigma-Aldrich). These NPC chemicals were added freshly before each media change and the final NKM is sterile filtered. The base medium N2B27 was composed of DMEM/F-12 (Thermo Fisher) and Neurobasal Medium (Thermo Fisher) at a 1:1 ratio, supplemented with 1:200 diluted N2 supplement (Thermo Fisher), 1:100 diluted B27 supplement without vitamin A (Thermo Fisher) and 1:100 diluted penicillin/streptomycin/glutamine (PSG, Thermo Fisher). Culture medium was exchanged daily and cells were split every 5–7 days when confluency of at least 80% was reached. At high cell numbers, the volume of cell culture medium was increased. Splitting was performed at a ratio of 1:12–1:20 with accutase (Sigma-Aldrich) treatment for approximately 5 min at 37°C to generate a single-cell suspension. Earlier passages were expanded in a lower splitting ratio of 1:8. Next, the NPC suspension was transferred to a falcon containing DMEM/F-12 with 0.1% bovine serum albumin (BSA, Thermo Fisher) to stop the enzymatic digestion and centrifuged at 300 g for 4 min. Finally, the cell pellet was resuspended in NKM and seeded out in fresh Matrigel-coated 6-well plates.

#### Differentiation of NPC into neurons

The differentiation protocolused here generates midbrain NPC-derived neurons. A day prior to the start of the differentiation, 300000 NPCs were plated onto a Poly-L-Ornithine (PLO)-coated well of a 12-well plate in NKM. The medium was changed after 24 h to Neuronal Induction Medium (NIM) to initiate the differentiation process of the NPC. Therefore, NIM contains 1 ^µ^M SAG, 75 ^µ^M Ascorbic Acid, and the neurotrophic growth factors BDNF and GDNF added each 2 ng/ml in addition to the base medium N2B27. This medium was exchanged daily for five days. On day 6 to day 9 Neuronal Differentiation Medium^+^ (NDM^+^) was used and medium change was performed every second day. On day 10 the Activin A was removed from the medium so that the cells were maintained in Neuronal Differentiation Medium (NDM) until the end of the differentiation on day 23. A schematic representation of the differentiation process from NPC to neurons is shown in Figure 18A. Cells were carefully re-plated with Accutase between day 8 and 12 depending on the experiment. An additional re-plating of neurons was performed by washing the cells with medium instead of PBS and carefully detaching them with TripLE (Gibco). After long-term culture on PLO the neuronal networks can detach from the well. Thus, NDM was aspirated with a manual pipette instead of vacuum when performing media changes.

#### Cell transfection

Hela cells were transfected with the reagent Polyethylenimine (PEI, Polysciences Inc.). PEI is a stable cationic polymer. For the transfection suspension, 2 ^µ^g plasmid DNA was mixed with 240 ^µ^l Opti-MEM medium followed by the addition of 4 ^µ^l PEI. The DNA:PEI transfection suspension was incubated for 1 h at room temperature and added dropwise to HeLa cells. After 18-24 h, the medium was exchanged - NPCs were transfected with TurboFectin 8.0 from OriGene. This transfection reagent is based on cationic lipids. Plasmid DNA was diluted in NKM without B27 and PSG and vortexed gently. TurboFectin 8.0 was added in a threefold greater volume than the DNA volume and carefully pipetted up and down. The transfection solution was incubated for 30 min at room temperature, then added dropwise to NPC. The transfection suspension was exchanged with new NKM after 24 h. The best transfection efficiency (≈ 0.001%) of NPC in a 12-well plate was found using 0.8 ^µ^g of total plasmid DNA.

#### Growth Curve

To create a growth curve, 35000 cells were seeded in 24-well plates. After adhesion a treatment was applied for 24 h. On the first day of measuring, the cells were washed with PBS and medium was exchanged. Cells in the first row of the plate were stained with Hoechst 33342 (0.1 ^µ^g/ml) and the fluorescent scan was performed at 37°C and 5% CO_2_. This cell counting procedure was performed with an automatic imaging reader (Cytation 5, Agilent) for four days in intervals of 24 hours post treatment.

#### Immunofluorescence

For immunofluorescence staining, cells were cultured on coverslip glasses or Ibidi-8-well chambers with glass bottom. Washing with PBS was carried out twice after each step. Prior to staining, cells were fixed with 4% paraformaldehyde (PFA) for 20 min at room temperature. Residual PFA activity was quenched by incubation in 0.1 M glycine for 10 min. Subsequently, cells were incubated in 0.1% Triton X-100 in PBS for 5 min for permeabilization and blocked in 5% goat serum in PBS for 30 min. The primary antibody of interest was diluted in 5% goat serum in PBS and the glass coverslip was flipped in order to incubate cells in a 20 ^µ^l drop of antibody solution on parafilm for 1 hour at room temperature or overnight at 4°C. Next, the secondary antibody was diluted 1:400 and added to cells for 1 hour at room temperature. Three additional washing steps with water were carried out to reduce background signal. A drop of mounting solution was applied to the surface of a microscope slide. The coverslip was slowly taped onto the slide and gently tilted into the mounting medium to avoid the formation of bubbles. Imaging was performed After at least 24 h at 4°C.

#### Cell Cycle Analysis

The DNA content in the cells was measured by propidium iodide staining of liquid fixed cells followed by flow cytometry analysis. NPC and HeLa cells were harvested and the cell pellet was resuspended in 500 ^µ^l ice-cold PBS. Subsequently, 4.5 ml ice-cold 70% ethanol (in ddH_2_O) was added dropwise to the cells while vortexing. Cells were stored at -20°C for at least 2 h. Next, cells were centrifuged at 2500 rpm for 5 min at 4°C to remove ethanol, washed with ice-cold PBS and centrifuged at 3200 rpm for 5 min. Cells were resuspended in 1 ml of ice-cold PBS, transferred to a 2 ml tube and recentrifuged. The cell pellet was resuspended in 500 ^µ^l of FxCycle PI/RNAse Solution, gently vortexed and incubated in the dark for 30 min at room temperature. Flow-Cytometry was performed with a 488 nm laser and analysis was performed using the CytExpert Software.

### Biochemical assays

#### Mdivi-1 inhibition assays

Mdivi-1 [Sigma-Aldrich M0133] was dissolved in DMSO at a concentration of 50 mM, sterile filtered and stored at -20 °C. Working concentrations of Mdivi-1 have been prepared in cell culture media for cellular treatment or according assay buffers for enzyme activity assays. Acute cellular treatment was performed by 24 h incubation of cells in media containing 50 µM Mdivi-1, long-term cellular treatment for one week in 10 µM Mdivi-1 respectively. Control experiments were performed with equal volumes of DMSO.

#### SDS-PAGE and blotting

Protein concentrations in cell lysates were determined by Bradford using ROTI Nanoquant reagent. 4x SDS sample buffer (0.83% SDS; 6.25 mM TRIS; 7.25% Glycerine; 0.0083% MercaptoEtOH; 0.01% Bromphenol blue) was added to the cell lysate and boiled at 95°C for 5 min. For SCAFI immune-blotting the sample had to be shaken at 40°C for 10 min after addition of 4x SDS sample buffer. Equal protein quantities (HeLa 50 ^µ^g, NPC 50 ^µ^g, neurons 40 ^µ^g) were loaded on a 4%/12% gel. The extracted proteins were separated by SDS-PAGE at 50 V until the samples passed through the stacking gel, then at 100 V for 2 h. The separated proteins were transferred onto polyvinylidene difluoride (PVDF) membranes by wet blotting at 25 V overnight.

#### Immunoblotting and membrane imaging

The dry membrane was incubated in methanol and washed for 5 min in water. Next, three washes were performed with TBST (TBS buffer, 1% Tween) for 10 min each. The membrane was then blocked for 1 h in a 10% milk solution. Then the primary antibody of interest was diluted in 1% milk in TBST and left overnight at 4°C with gentle rotation. On the following day, the membrane was washed three times with TBST, then incubated for 1 h in the secondary antibody diluted in 1% Milk-TBST solution. Three washing steps with TBST followed by one washing step in Milli-Q water for 5 min each were performed. For detection, an enhanced chemiluminescent substrate (ECL) consisting of a 1:1 mix of Luminol/Enhancer & Peroxide solution was added to the membrane and incubated for 2-5 min. Finally, band signal detection was conducted with the ChemiDOC Infrared Imaging System (BioRad). The detection of phosphorylation of proteins as well as specific antibodies (for OPA-1, SCAFI, TIMBIM5 and SYP) required 5% BSA for blocking and 1% BSA for antibody dilution.

#### Isolation of Mitochondria

For isolation of mitochondria from cultured mammalian cells, the cells of two confluent T175 flasks were trypsinized, harvested with a cell scraper and cell pellets were frozen at -80°C. All following procedures were performed on ice. The cell pellet was briefly thawed and recentrifuged at 300×g for 5 min at 4°C. The supernatant was discarded and the pellet was resuspended in mitochondria isolation buffer (MIB) (0.32 M sucrose, 1 mM EDTA, and 10 mM Tris–HCl, pH 7.4). Next, the cell suspension was transferred into a Dounce homogenizer (B.Braun) with a tight fitted pestle to lyse cells. Mechanical homogenization was performed and the cell lysates were centrifuged at 800×g for 5 min at 4°C. This step was repeated until no cell pellet was visible any more. Finally, the supernatant was centrifuged at 10000×g for 10 min at 4°C followed by resuspension of the pellet in mitochondrial isolation buffer and concentration quantification.

#### BN-PAGE

Mitochondrial membrane protein complexes were separated using native gel PAGE as described in (Wittig et al. 2006). All steps were performed on ice. First, mitochondria pellets were obtained by centrifuging for 10 min at 9000×g. The pellets were resuspended in the blue native sample buffer (for 50 ^µ^g crude mitochondria: 5 ^µ^L NativePAGE sample buffer (2×), 8 ^µ^L 5% digitonin, and 7 ^µ^L water). Next, the detergent was added. Digitonin was used in a 6 g/g ratio (detergent/protein). After 10 min solubilization with the digitonin, the suspension was centrifuged for 20 min at 20000×g. The supernatant contains mitochondrial membrane proteins. 20% Glycerol and 25% of detergent concentration with Coomassie were added before lo^4^ading samples into the gel. 50 ^µ^g protein was loaded per lane and separated on a 3–13% acrylamide gradient gel. The power supply was set to 100 V for the first 60 min and then increased by 20 V in intervals of 1 h to 240 V until the desired separation was reached. After 4-6 h running time, the native gel was washed with ddH_2_O and stained with Simply Blue stain to visualize the native marker and total protein lanes. Next, the gel was electro-blotted onto Hybond-P-polyvinylidene fluoride (PVDF) membranes (GE Healthcare) using a wet blotting system at 25 V overnight. After finishing the transfer, the PVDF membrane was incubated for 15 min in a fixative solution (8% acetic acid) with gentle shaking, followed by three washing steps with ddH_2_O. After that, the membrane was air-dried for 30 min at room temperature. The membrane was either used directly for immunoblotting or stored covered in Whatman paper in the fridge overnight or at -20°C for extended storage. Coomassie staining of the full protein bands is printed on the membrane. First, the native marker bands were marked with a pen then the staining was removed through 5 min washing in methanol with gentle shaking. Finally, the usual western blot protocol was performed.

#### Complex I In-Gel Activity Assay (CI-IGA)

To analyze complex I activity, the blue native gel was incubated for 24 h in 20 ml complex I substrate solution until violets bands were clearly visible indicating active complex I. The reaction was stopped by denaturing the native complexes with 10% acetic acid solution. Finally, the gel was washed with water and a picture was taken.

#### Enzyme activity of isolated OXPHOS complexes by spectrophotometry

The individual respiratory complex activity assays were adapted from Andrea Curtabbi. Isolated mitochondria were frozen at -80°C, thawed on ice, resuspended briefly and refrozen. This freeze-thaw cycle was repeated three times to disrupt the mitochondrial membrane. Repeated freeze-thaw cycles in hypotonic buffers increase enzymatic activity and sensitivity to Complex I inhibitors such as Rotenone (Spinazzi, Casarin et al. 2012).

A base buffer for individual complex enzyme activity (BICA) (250 mM Sucrose, 10 mM Tris/MOPS, 1 mM EGTA) was prepared. For each assay, 500 ^µ^l of the according assay buffer, depending on the type of Complex I enzyme assay (NADH:DQ or NADH:FeCy), was pipetted into wells of a 96-well plate with flat bottom. Next, 20 ^µ^g of isolated mitochondria was added per well and mixed without formation of bubbles. Finally, the Cytation 5 was set up in a kinetic readout mode to determine the appropriate optical density (OD) in 12 s intervals. All spectrophotometric assays are time-dependent and need an electron donor as well as an electron acceptor to investigate a concrete respiratory chain complex activity. A blank control containing assay buffer without mitochondria was always recorded in parallel for background subtraction.

#### NADH:DQ activity assay

To measure NADH oxidation of isolated complex I via the 6-decyl derivative of ubiquinone (DQ) as electron acceptor, the further electron transfer from Complex I to Complex III had to be blocked by inhibition of Complex IIII with Antimycin A. The spectrophotometric measurement of NADH:DQ oxidation at OD 340 was performed in NADH:DQ activity buffer (130 µM NADH, 130 µM DQ, 4µM Antimycin A in BICA) at 35°C for 5 min.

#### NADH:FeCy activity assay

NADH dehydrogenases catalyze rapid reduction of ferricyanide (FeCy) by NADH. The NADH:FeCy oxidation is not sensitive to Rotenone and probably FMN is the reactive site donating electrons to FeCy (Vinogradov 1998). The spectrophotometric measurement of NADH:FeCy oxidation at OD 340 was performed in NADH:FeCy activity buffer (130 µM NADH, 1 mM FeCy, 4µM Antimycin A in BICA) at 32°C for 8 min.

#### Determination of gene expression via quantitative PCR (qPCR)

Total RNA of cell pellets was obtained with the Monarch Total RNA Miniprep Kit (NEB #T2010S), and cDNA was transcribed with the RevertAid First Strand cDNA Synthesis Kit (Thermo Scientific #K1621). Quantitative PCR (qPCR) was performed on the StepOnePlus Real-Time PCR System (Applied Biosystems). The PCR reaction was prepared from PowerUP SYBR Green Master Mix (Applied Biosystems # A25741), 50 ng cDNA, and 0.1 nM of each forward and reverse primer per sample. The PCR protocol started with an initial denaturation step of 10 min, followed by 40 cycles of 95°C for 15 s and 60°C for 60 s. The PCR cycle numbers at which the reaction curve intersects the threshold lines (C*_T_* values) were exported by the StepOne Software 2.3 and the relative gene expression (RQ) was calculated according to:

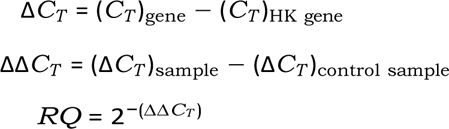

where *β*-V Tubulin was used as a housekeeping gene and Δ*C_T_* was normalized to a DMSO control of the according cell type. Normalization of Δ*C_T_* on NPC wt was performed when the relative gene expression of multiple samples was investigated to observe changes throughout the differentiation process. All used primers were synthesized by Eurofins Genomics. Quality control for primer accuracy and efficiency was performed manually for each pair of Primers via Relative Standard curves in a 1:5 serial dilution of 200 ng cDNA.

#### Oxygen consumption measurements

All measurements of oxygen consumption rates (OCR) and extracellular acidification rates (ECAR) were performed with a Seahorse XF96 Extracellular Flux analyzer (Agilent) in a time-dependent experiment on cultured HeLa cells or differentiated NPC-derived neurons. For measurements of the mitochondrial respiration, the cells were seeded in a Seahorse XF Cell Culture Microplate (96 wells) one the day before of the assay (35000 cells/well). To hydrate the sensor cartridge 200 ^µ^l XF Calibrant (pH 7.4) was added to each well and incubated at 37°C and environmental CO_2_ overnight. On the day of performing the assay, the cells were incubated in 180 ^µ^l assay medium (25 mM D-Glucose or galactose, 1 mM pyruvate, 2 mM L-Glutamine in XF Base Medium Minimal DMEM 0 mM glucose, pH 7.4) for 1 h at 37°C and environmental CO_2_.

#### Mito-stress test

For the so-called mito-stress test, the following compounds were added subsequently: oligomycin, an inhibitor of ATP synthase; Carbonylcyanid-p-trifluoromethoxyphenylhydrazon (FCCP), an uncoupler; Rotenone, an inhibitor of Complex III, plus Antimycin A (AA), an inhibitor of complex IV.

Inhibitor stock solutions were prepared with assay medium and added to the ports of the sensor cartridge. The concentrations of inhibitors in the ports were tenfold higher than the desired final concentration for the cells, because of the dilution factor created by injecting the inhibitors into the assay medium. Final well concentrations were 1 ^µ^M for oligomycin, 0.5 ^µ^M (0.75 ^µ^M for neurons) for FCCP and 1 ^µ^M for Rotenone/antimycin A. The measurements were performed at 37°C. With the final injection, the cells were stained with Hoechst 33342 (2 ^µ^g/ml) to allow for counting the cell number after finishing the assay (conducted with the Cytation 5). This allows for the normalization of the mitochondrial respiration to the cell number.

#### Respiratory complex activities

Activities of individual respiratory chain complexes can be analyzed after permeabilization of membranes with digitonin and application of specific OXPHOS substrates. 1x MAS buffer is prepared from 3x MAS buffer (660 mM mannitol, 210 mM sucrose, 30 mM KH_2_PO_4_, 6 mM HEPES, 3 mM, EGTA, 15 mM MgCl_2_) with the following supplements: 10 mM glutamate, 10 mM malate, 0.5 µM FCCP and 0.0006% digitonin. This buffer is referred to as MAS++. The prewarmed MAS++ buffer (37°C) is adjusted to pH 7.4 (with 0.1 M HCl or 0.1 M KOH). After calibration, the growth medium (glucose medium) is exchanged by MAS++ (180 µl per well; after washing 1x with 60 µl MAS++ per well).

The substrates for CI (glutamate and malate) and for CII (succinate and G-3-P) were applied to measure in CI/CIII/CIV- or CII/CIII/CIV-driven oxygen consumption. Rotenone and antimycin A were added to block CI and CIII which led to a decrease of oxygen consumption. Tetramethyl-p-phenylenediamine (TMPD) and ascorbate were injected for determination of CIV-dependent oxygen consumption alone. The 1x MAS^++^ buffer with the substrates for complex I was added prior to the assay. The other substrates and inhibitors were added during the assay in a specific order to measure the respiratory activities of single complexes. For determination of the respiratory complex activities, 35000 cells cells/well were seeded in a Seahorse XF Cell Culture Microplate (96 wells) on the day before of the assay. To hydrate the sensor cartridge 200 ^µ^l XF Calibrant (pH 7.4) was added to each well and incubated at 37°C and environmental CO_2._ On the day of experiment, the substrates (10 mM succinate, 5 mM G-3-P (glycerol-3-phosphate, 10 mM ascorbate and 100 µM TMPD) were prepared with 1x MAS^++^ buffer, as well as the inhibitors Rotenone (final concentration 0.5 µM) and antimycin A (final 0.5 µM) and then added to the ports of the sensor cartridge. Measurements are performed at 37°C. With the final injection, the cells were stained with Hoechst 33342 (2 ^µ^g/ml) to allow for counting the cell number after finishing the assay (conducted with the Cytation 5). This allows for the normalization of the mitochondrial respiration to the cell number.

#### Microelectrode array (MEA) measurements

Electrophysiological characterization and neuronal activity of neurons were performed by using 9-well on microelectrode array (MEA) chips (USB-MEA256system, Multichannel Systems) as previously described (Renner, Grabos et al. 2020). The MEA chips were plasma-cleaned and coated with a 1:75 Matrigel (Corning) dilution in KO-DMEM (Gibco) and 5% KSR (Thermo Fisher) overnight and additionally incubated with FBS (Sigma-Aldrich) at 37°C for 30 min. Cell clusters were detached by using Try-pLE (Gibco) transferred to the electrode area of the MEA chips and attached for about 96 h. The measurements of the cells were performed at a temperature of 37°C. Electric field potentials were first recorded under basal conditions without any activator or inhibitor to record a reference signal. To identify neurotransmitter responsive network activity, different pharmacological agonistic and antagonistic modulators were applied to each sample chamber of a MEA chip and electric field potential was recorded. The activators glutamate/glycine (100 ^µ^M each, Sigma-Aldrich), dopamine hydrochloride (10 ^µ^M, Sigma-Aldrich) and GABA (10 ^µ^M, Sigma-Aldrich) were applied to selectively detect neurotransmitter responsive neuronal networks. After recording the activator responsive signals, selective inhibitors for the respective pathways were applied to the same well. The used inhibitors were risperidone (10 nM, Sigma-Aldrich), bicuculline (1 ^µ^M, Sigma-Aldrich) and ketamine (10 ^µ^M, Sigma-Aldrich). For recording the datasets, Cardio2D software (Multi Channel Systems MCS GmbH, Reutlingen, Germany) was used. Data was analyzed using the softwares Cardio2D+ (Multi Channel Systems MCS GmbH, Reutlingen, Germany) and Origin v9.0 (OriginLab Corporation, Northampton, MA, USA). MEA analysis was performed on n=3 independent replicates.

### Imaging

For imaging, a cLSM (TCS SP8 SMD) equipped with a 60x objective (NA 1.35, N/0.17/FN26.5, Leica), two Hybrid GaSP-detectors (HyD) and a tunable white-light laser was used. The device was also equipped with a time correlated single photon counting (TCSPC) set-up for fluorescence lifetime imaging microscopy (FLIM). Live cell imaging was performed at 37°C and 5% CO_2._

To monitor supercomplex formation via FLIM, 30000 HeLa cells were seeded into an ibidi-8-well and transfected with Cox8a sEcGFP. For excitation, the 488 nm wavelength of a white light laser was used with a repetition rate set to 40 MHz. Fluorescence signals were detected by a Hybriddetector (Leica), emission range was adapted to an avalanche diode with an emission filter FF1 520/35 (Semrock). Photon traces were recorded with TCSPC, which allowed time gating. Data acquisition was stopped when at least 1000 photons for the brightest pixel were collected.

Data from TCSPC experiments was analyzed with SymphoTime software PicoQuant. Fluorescence decay curves from ROIs displaying at least 80% of the mitochondrial network were fitted via tailfit of a bi-exponential decay function (including/substracting the Instrument Response Function = IRF). The average lifetime *τ*_Amp_ abbreviated with *τ* was used for evaluation. The resulting values for the fluorescence lifetime were displayed on a false color scale.

#### Mitochondrial mass and morphology

MitoTracker™Green FM (MTG) is a membrane potential-independent mitochondrial tracker with excitation/emission maxima ∼490/516 nm, which accumulates in mitochondria and binds covalently to mitochondrial proteins by reacting with free thiol groups of cysteine residues. However, a minimum amount of membrane potential is needed to allow the incorporation of the dye. MTG (Invitrogen, #M7514) was used to stain mitochondrial mass for Mitochondrial Network analysis and normalization of dyes monitoring specific bioenergetic parameters. Cells were stained at a final concentration of 100 nM for 30 min at 37°C and 5% CO_2_. Next, one washing step with PBS and two washing steps with culture medium were performed on HeLa cells to remove cytosolic background signal. For neurons, staining medium was aspirated carefully and three washing steps with NDM were performed prior to imaging with the cLSM.

#### Mitochondrial membrane potential

Tetramethylrhodamine ethyl ester (TMRE) is a cell-permeant, cationic, fluorescent dye with excitation/emission maxima ∼549/575 nm, and was used to measure mitochondrial membrane potential *ψ_m_*. The dye accumulates in the mitochondrial matrix and high TMRE signal indicates a higher ΔΨ_m_, when the dye is used in the non-quenching mode. Cells were stained with 7 nM TMRE for 30 min at 37°C and 5% CO_2_ to measure mitochondrial membrane potential. Cells were simultaneously stained with MTG for normalization on the mitochondrial mass. Next, washing steps were performed as previously described for MTG staining and fresh imaging medium containing TMRE without MTG was added. Cells were imaged after an equilibration time of 10 min at 37°C and 5% CO_2_. After imaging, 1 ^µ^M FCCP was added to the medium, to determine how the TMRE signal fades away over in a time period of 30 min due to uncoupling.

#### Mitochondrial ATP levels

ATP-Red Live cell dye (Biotracker) reports mitochondrial ATP levels, when a negatively charged ATP breaks the covalent bonds between boron and ribose, causing ring-opening and fluorescence. The red fluorescent dye has excitation/emission maxima of ∼510/570 nm. Cells were incubated with 5 ^µ^M ATP-Red for 15 min at 37°C. Cells were simultaneously stained with MTG for normalization on the mitochondrial mass. Next, washing steps were performed as previously described for MTG staining and fresh medium was added prior to imaging.

#### Mitochondrial superoxide levels

MitoSox (Thermo Scientific) detects superoxide localized in the mitochondria. Cells were stained with 2.5 ^µ^M MitoSox for 30 min at 37°C and then washed three times with medium before imaging. As a control for lipid peroxidation, a sample was treated with 300 ^µ^M Tert-Butyl Hydroperoxide (TBH70X, Luperox) for 30 min.

#### Mitochondrial lipid peroxidation

MitoCLox is a novel dye, that determines Lipid Peroxidation in response to oxidative stress (Lyamzaev, Panteleeva et al. 2020). Cells were stained with 200 nM MitoCLox for 2 h and subsequently washed 3 times with medium. Imaging of cells was performed recording z-stacks. MitoCLox emission was recorded at 488 nm and 559 nm.

#### Calcium imaging

Yellow Chamelaeon (YC3.6) is a biosensor that is sensitive to cytosolic Calcium ([Ca^2+^]_cyto_). The sensing mechanism of Ca^2+^ is based on the CaM binding peptide between the fluorophores YFP and CFP allowing for fluorescence resonance energy transfer (FRET) when Calcium is bound. The intensity of its FRET signal changes as the Ca^2+^ level within the cell rises and falls. The emission ratio of the fluorophores YFP/CFP indicates the amount of bound calcium. Cytosolic uptake was measured by ratio metric imaging of YC 3.6-transfected HeLa cells via stimulation with 2 ^µ^M CaCl_2_ for 8 min after recording a baseline of 2 min. Mitochondrial Calcium ([Ca^2+^]_mito_) levels were measured with the Chameleon biosensor mt4D3cpv, which contains a targeting sequence for the mitochondrial matrix and a circular permuted Venus as an acceptor fluorophore for FRET.

Fura-2 AM, is a membrane-permeant derivative of the fluorescent dye Fura-2. Cytosolic Calcium concentrations were measured by ratiometric imaging. When added to cells, Fura-2 AM crosses cell membranes and once inside the cell, the acetoxymethyl groups are removed by cellular esterases. Removal of the acetoxymethyl esters regenerates Fura-2, the pentacarboxylate calcium indicator. Measurement of Ca^2+^-induced fluorescence at both 340 nm and 380 nm allows for calculation of calcium concentrations based on F340/380 ratios. 0.05% Pluronic was added to increases the incorporation of Fura-2 AM into the cells. Cells were stained with 5 ^µ^M Fura-2 AM and calcium transients were recorded with a Nikon Eclipse Ti-S microscope equipped with a 40X oil immersion objective with 0.5 frames per second in the MetaFluor Fluorsecence Ratio Imaging Software. Basal and stimulated neuronal activity was determined by cellular calcium uptake pre and post activation with glutamate/glycine (each 100^µ^M). Measurement of neuronal calcium uptake was quantified by the area under curve (AUC) starting from the baseline.

### Image processing

#### Mitochondrial Network analysis (MiNa)

To quantify Mitochondrial morphology, images of MTG stained HeLa cells were analyzed with the mitochondrial network analysis (MiNa) macro in ImageJ. For the analysis, a binary image is produced of the mitochondrial signal and skeletonized to remove non-mitochondrial external pixels (Valente, Maddalena et al. 2017). Next, the image is segmented into four morphological categories: networks, large/round, rods and punctate. The extent of fragmentation into small, punctate or linear structures versus the extent of larger and more branched structures was determined by the MiNa algorithm. The morphological analysis by MiNa was restricted to the mean branch length and mitochondrial mass in this thesis. Mean branch length indicates the average length of all branches (the distances between connected end point or junction pixels) and mitochondrial footprint is the mitochondrial mass of the total area.

#### *In silico* Molecular Docking experiments with complex I

All structural alignments and docking experiments were performed by using YASARA structure 2023 via implemented and adapted macros. The previously published inhibitor-bound complex I cryo-EM structure obtained from *Mus musculus* with PDB number 6ZTQ (Bridges, Fedor et al. 2020) was used as a docking platform after removing the inhibitor piercidin A. Complex I from *Bos taurus* (PBD: 5LDW) (Zhu, Vinothkumar et al. 2016) and from *Homo sapiens* (PBD: 5XTD) (Guo, Zong et al. 2017) were used for further experiments with respiratory complexes of other hosts. Prior to docking, the four ligand structures of rotenone, Mdivi-1, Q1 and Q10 were imported as SMILES strings in YASARA and energy minimized. Next, the previously cropped piercidin A was added to the ligands as a control. Global docking was performed using the previously described cryo-EM structures with a simulation box surrounding the Q-cavity of complex I as binding site and the modified YASARA macro dock run.mcr with 100 runs per ligand. Additionally, cryo-EM structures were cropped around the binding domain for docking experiments with AutoDock. For subsequent analysis of interacting residues, the docking conformation with highest binding energy was chosen for each ligand and compared to the interactions in the inhibitor-bound cryo-EM structure. N=4 docking processes with n=25 docking poses each were performed. According to recent publications (Bridges, Fedor et al. 2020) (Murai 2020) and specific residues were identified as crucial for ligand interactions in the Q-cavity and therefore focused on during analysis.

### Statistical analysis

All graphs were generated with Origin Pro. Pooled data of biological replicates were checked for significant outliers via Gubbs test. After removal of significant outliers, statistical analysis for normally distributed data was performed via one-way ANOVA with Tukey post-hoc tests, while not normally distributed data was analyzed via Kruskal-Wallis-ANOVA with Dunn’s post-hoc tests. Asterisks indicate level of statistical significance: * *p* ≤ 0.05, ** *p* ≤ 0.01, \*\*\**p* ≤ 0.001, tendency *p* ≤ 0.1 or not significant *p >* 0.05. *N* indicates the number of independent experiments performed with a different batch of cells or for differentiated neurons a new differentiation and *n* represents the number of total measurements.

## Acknowledgments

We thank Frank Schmelter for valuable technical assistance, Silke Morris for proof reading the manuscript and Andreas Curtabbi for inspiring discussions and suggestions as well as technical support. We thank Isabel Aymanns who contributed important supplementary data (Calcium imaging in neurons). The project was funded by the GRC (INST 190/167-2; SFB944). Microscopes of the Münster Imaging Network were used.

## Authors contributions

NM and KBB designed the experiments, validated the data. NM performed the experiments. NR, PD and GS contributed to the methodology. KBB contributed to resources; KBB and NM contributed to writing— original draft preparation; KBB and NM visualized the study; KBB and NM supervised the data; KBB contributed to project administration; KBB acquired funding. All authors have read and agreed to the published version of the manuscript.

## Materials availability

This study did not generate new unique reagents nor genetically modified organisms.

