## supplementary Figures for "Mdivi-1 affects neuronal activity by inhibition of Complex I and respiratory supercomplex assembly"

### Supplementary information

#### Differentiation of neurons from neuronal progenitor cells

The NPC were a gift from Thomas Gasser, Universitätsklinikum Tübingen, Germany. The differentiation and maturation of NPC to neurons was conducted following usual protocol (Figure S 1A). Maturation was controlled by the gene expression profile of markers for pluripotency respectively neurons in culture samples taken in intervals of 5-6 days. The mRNA expression of the pluripotency markers SRY (sex determining region Y)-box 2 (Sox-2), octamer-binding transcription factor 4 (Oct-4) and Krüppel-like Factor 2 (KFL-2) were significantly decreased in samples from day 23 (neurons) compared to day 0 (NPC) of differentiation (Figure S1B). Furthermore, the genes encoding for the Microtubule-associated protein 2 (MAP2) and the Growth Associated Protein 43 (GAP43) showed an increasing trend for mRNA expression throughout the differentiation process, resulting in a significant upregulation of these genes in neurons compared to NPC. An immunofluorescence co-staining of MAP2 and the neuron-specific class III beta-tubulin (TUJ-1) showed numerous neurons in differentiated cell culture (Figure S1C), which goes in line with the increase of MAP2 gene expression. Additionally, a 1:4000 IF staining of acetylated Tubulin indicated specification of TUJ-1$_+$ cells into axonal sub-compartments of neurons (Figure S1D), which are required for functional neurons. To verify the functionality of the cell model, differentiated neurons were electrophysiologically characterized by multi-electrode array (MEA) measurements. To identify neurotransmitter responsive network activity, different pharmacological agonistic and antagonistic modulators were applied as shown in Figure S1G. The neuronal activity determined by these measurements revealed a mixed neuronal culture, which was primary responsive to glutamate/glycine and dopamine. The presence of neuronal subtypes in the culture were shown by immunofluorescence co-staining of neurons with Tyrosinhydroxylase (TH) and choline-acetyl transferase (CHAT). Tyrosinhydroxylase is an important enzyme catalyzing L-DOPA production, which is an intermediate required for dopamine synthesis. Figure S1E shows a subpopulation of TH$_+$ neurons, indicating the presence of dopaminergic neurons in the culture. Staining of CHAT revealed neurons, that contain the transferase enzymes for the synthesis of the neurotransmitter Acetyl Choline, indicating the presence of cholinergic neurons (Figure S6A).

#### Mdivi-1 increases the mitochondrial mass in HeLa cells

To investigate if Mdivi-1 affects the mitochondrial morphology and network, HeLa cells were stained with Mito Tracker Green FM (MTG) after treatment with 10 µM Mdivi-1 for 24 h or 50 µM Mdivi-1 for 24 h (Figure S2). As controls, in one sample, complex I was inhibited by Rotenone for 24 h (1 µM) and one cell sample was transfected with Drp1K38A, which is fission-incompetent (Karbowski, Lee et al. 2002). Image analysis of single cells was performed with the MiNa (Mitochondrial Network analysis) macro in ImageJ to investigate changes in mean branch length (average length of all branches) and mitochondrial footprint (mitochondrial mass). Cells treated with increasing concentrations of Mdivi-1 did not alter the mean branch length of the mitochondrial network compared to DMSO treated cells (Figure S2B). Additionally, two replicates contained cells transfected with Drp1K38A mtEGFP (the dominant negative inactive form of Drp1, tagged with an EGFP) as a control for impaired mitochondrial fission. Mito Tracker Deep Red FM (MTDR) was used to stain and quantify these samples. HeLa Drp1K38A mtEGFP cells had a median branch length of 1.28 µm ± 0.18 µm; SD), which is 78% greater than the median branch length of DMSO treated cells, indicating hyperfused mitochondria. The mitochondrial footprint of cells treated with 50 µM Mdivi-1 was significantly increased. Furthermore, the mean branch length and the mitochondrial footprint of cells treated with the CI inhibitor Rotenone was decreased, indicating mitochondrial fragmentation. The protein level of the voltage-dependent anion-selective channel (VDAC) of the outer mitochondrial membrane was increased in acute Mdivi-1 treated HeLa cells and neurons, which also indicates an elevated mitochondrial mass (Figure S2C). Accordingly, the relative mRNA expression of mitochondrial transcription factor A (TFAM) shown in Figure S2D was increased in acute Mdivi-1 treated HeLa cells. Together this data shows an increase in mitochondrial mass of HeLa cells and neurons.

### Supplementary Figures

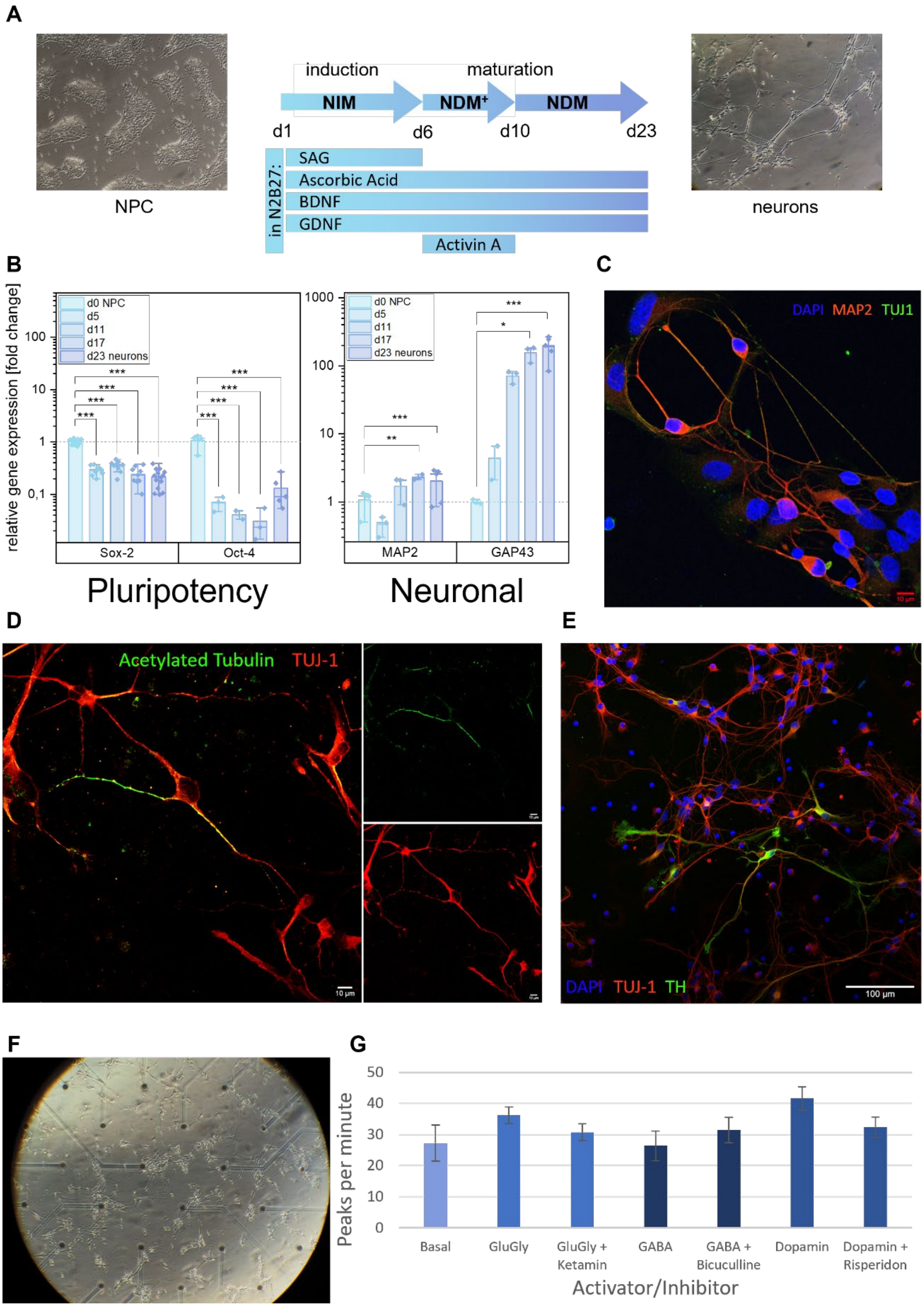


**Figure S 1: Generation of functional NPC-derived midbrain neurons.** (A) Schematic overview of NPC differentiation process with exemplary cell culture images. (B) Relative gene expression of markers for pluripotency (Sox-2 and Oct-4) decrease, while markers for neurons (MAP2 and GAP43) increase during the differentiation process (N=1, n=9). (C,D,E) Specific neuronal immunofluoresecence stainings of NPC-derived neurons. (F) Exemplary image of neurons post transferring on a multi electrode array chip. (G) Electrophysiological characterization of neuronal culture (N=3, n=9).

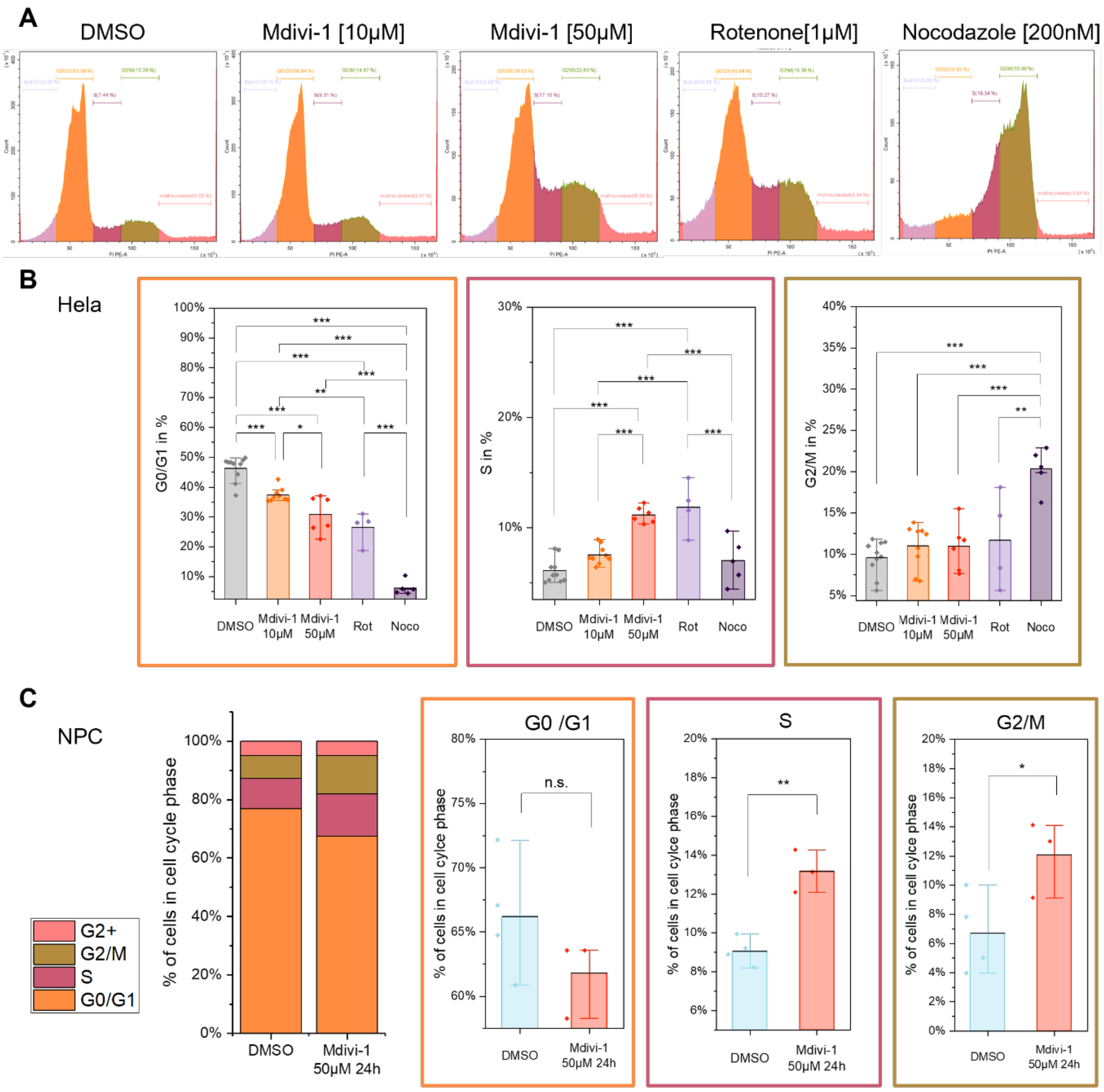


**Figure S 2: Mdivi-1 impairs cell growth due to stochastic failure of mitosis.** (A) Exemplary flow-cytometry results of HeLa cells stained with Propidium Iodide/RNA. (B,C) quantification of cell cycle analysis reveals accumulation of cells in S and G2/M phase in (B) HeLa cells (N=4-10, each n=200000, ANOVA, KW) and (C) Neural Progenitor cells (N=4, each n=200000, KW, ANOVA). p

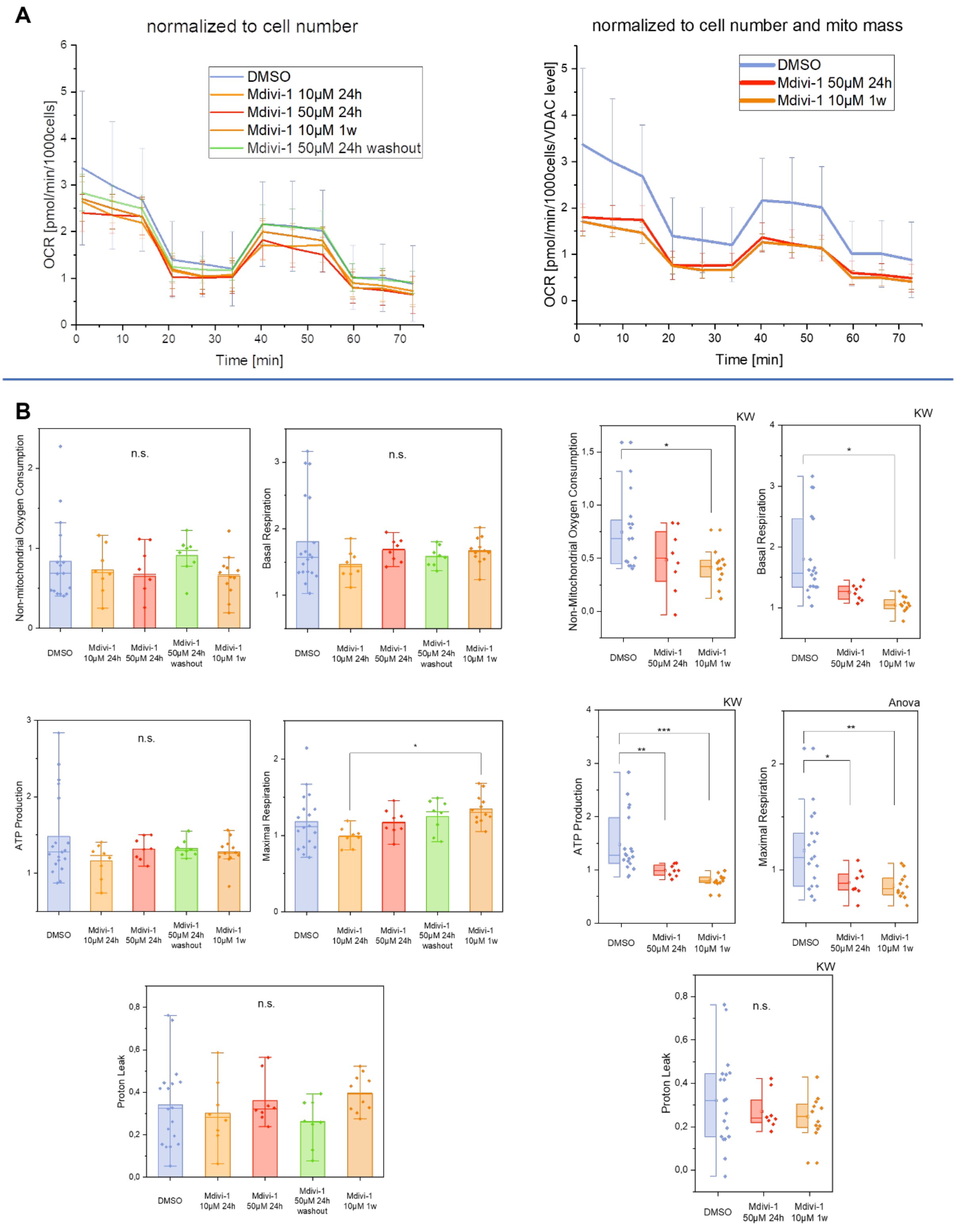


**Figure S 3: Effects of Mdivi-1 on mitochondrial respiration in neurons.** (A) Mito Stress test to determine OCR in neurons, inhibitors for ATP synthase (oligomycin¸ 1 µM), uncoupler trifluoromethoxy carbonylcyanide phenylhydrazone (FCCP, 1 µM) and inhibitors for complex I (Rot., rotenone; 0.75 µM) and complex III (AA; antimycin A; 1 µM) were subsequently added. Oxygen consumption rates (OCR) were determined with an automatic flux analyzer (Seahorse XF96/Agilent) and normalized to cell numbers (left panel) or mitochondrial mass (right panel). The normalization on the mitochondrial mass was done by using the factor that describes the mean protein level of VDAC normalized to TUJ-1. (B) Mitochondrial basal, ATP production-related and maximal respiration, non-mitochondrial oxygen consumption and proton leak in living cells (control and Mdivi-1 treated) were determined by OCR [pmol/min/1000cells] or containing additional normalization on the mitochondrial mass. (N=1, n>=8, left panel all tested with KW, right panel statistic test indicated).

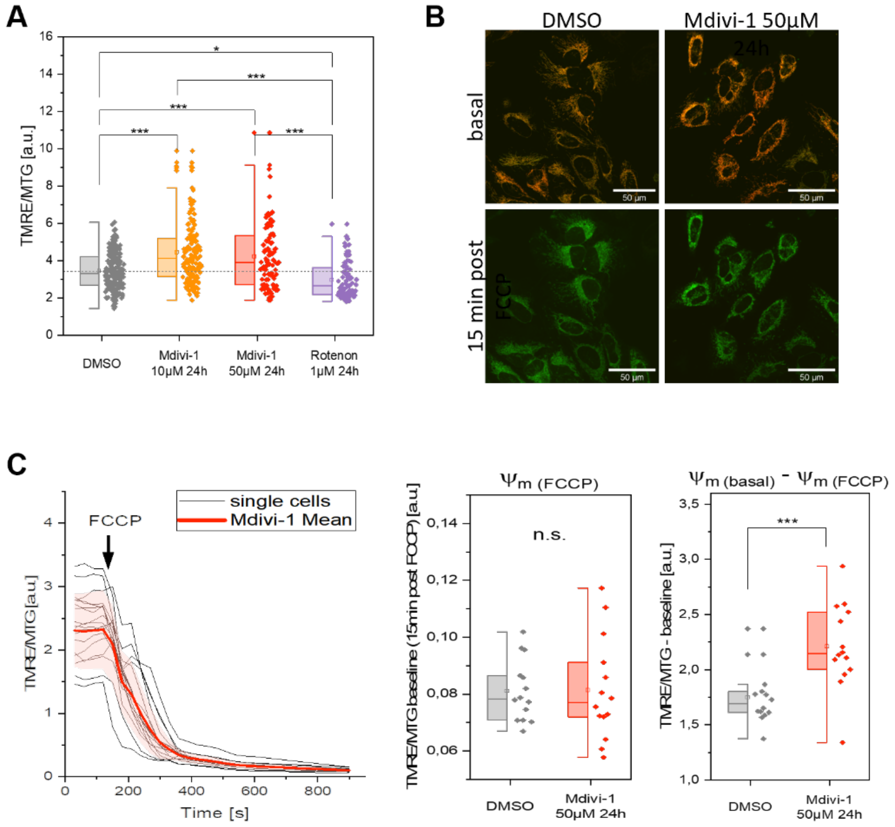


**Figure S 4: Mdivi-1 treated HeLa cells display an increased mitochondrial membrane potential.** (A) Effect of Mdivi-1 on mitochondrial membrane potential in HeLa cells (N=3, $n_{DMSO}$=200, $n_{Mdivi\text{-}1\ 10\ \mu M\ 24h}$=138, $n_{Mdivi\text{-}1\ 50\ \mu M\ 24h}$=94, $n_{Rotenone\ 1\ \mu M\ 24h}$=92). (B) $\Delta\Psi_m$ before and after FCCP-decoupling. Control and Mdivi-1-treated HeLa cells were stained with TMRE and MTG, imaged and exposed to FCCP (1 µM) for 15 min. (B) Decay of $\Delta\Psi_m$ in single cells due to FCCP treatment. (C) Quantification of $\Delta\Delta\Psi_m$. The $\Delta\Psi_m$ baseline 15 min post FCCP addition is unchanged, while the resting $\Delta\Psi_m$ is significantly increased (N=1, n=15 per condition, ANOVA).

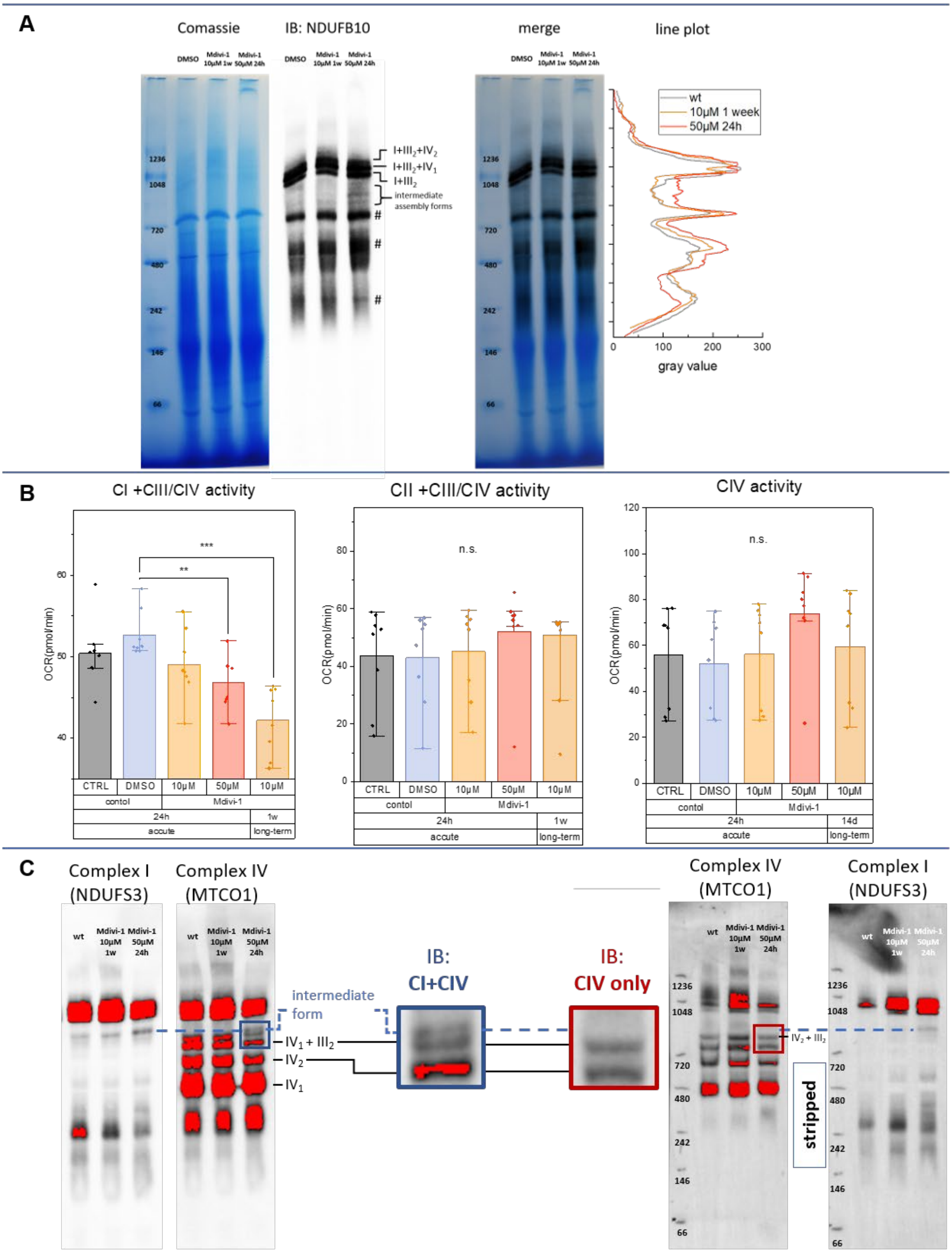


**Figure S 5: Mdivi-1 affects CI-dependent SC activity.** (A) Blue Native gel with isolated mitochondria of Mdivi-1 treated HeLa cells and immunoblotting of NDUFB10 shows an increased intermediate CI assembly form (N=1). (B) Determination of respiratory complex activities via Seahorse in neurons reveal a decrease in Complex I dependent respiration and no alterations for Complex II and IV dependent respiration (N=1, n=8). (C) Representative inverse immunoblotting with antibodies against NDUFS3 (Complex I) and MTCO1 (Complex IV). Overexposure visualizes that the intermediate Complex I assembly form does not contain Complex IV.

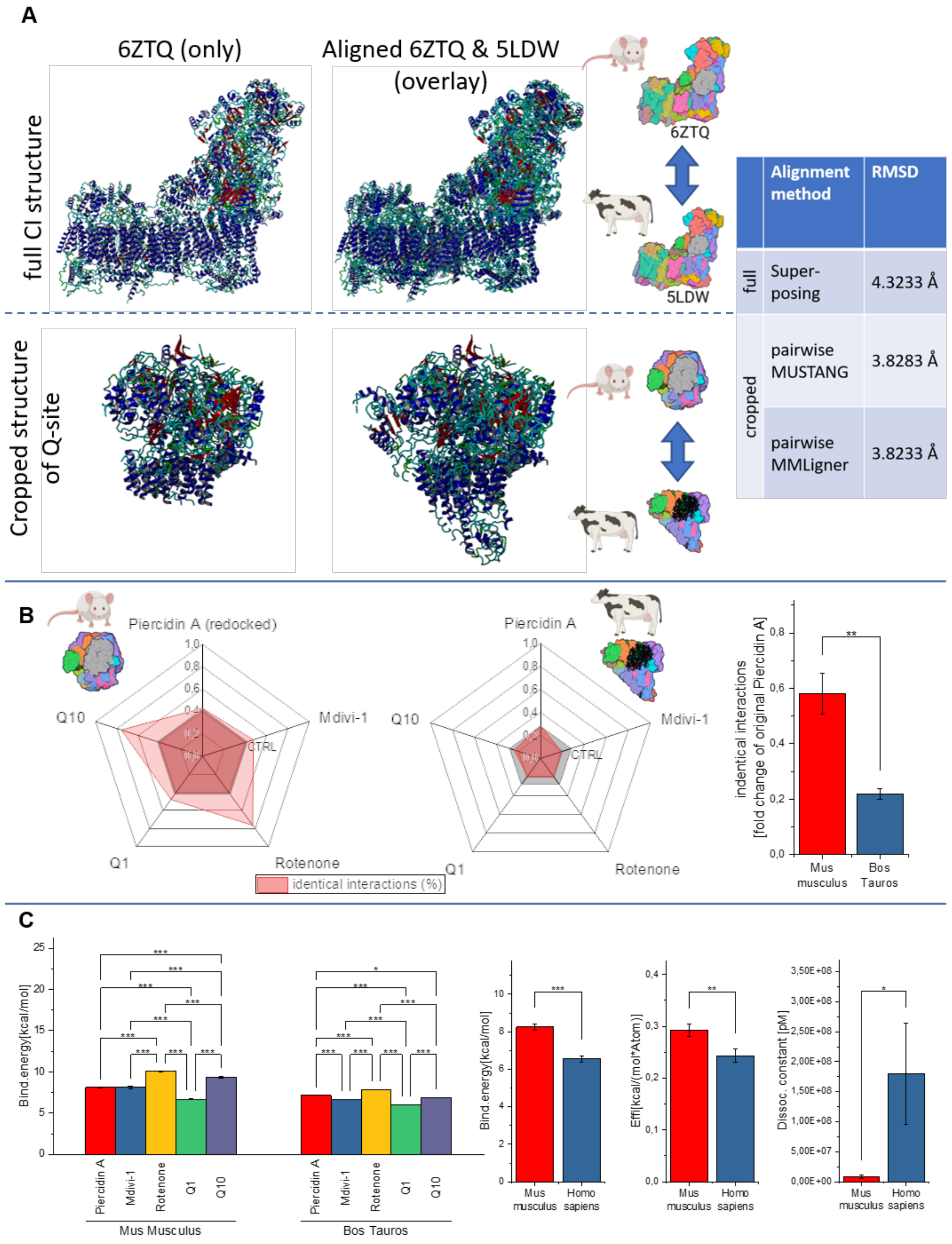


| | Alignment method | RMSD |
|---|---|---|
| full | Super-posing | 4.3233 Å |
| cropped | pairwise MUSTANG | 3.8283 Å |
| | pairwise MMLigner | 3.8233 Å |

**Figure S 6: Supplementary Results for molecular docking.** (A) Alignment of murine and bovine Complex I structures. (B) Quantification of identical interactions between bovine Complex I and binders piericidin A, rotenone, Q1, Q10 and Mdivi-1 shows less interactions compared to the redocked piercidin A in murine Complex I (left=comparison of individual best poses generated by Autodock, right=pooled poses of all ligands, n=5 poses per structure). (C) Respective binding energy of best docking poses (n=5 per condition) and parameters of pooled docking poses (N=4 docking processes, n=25 docking poses).

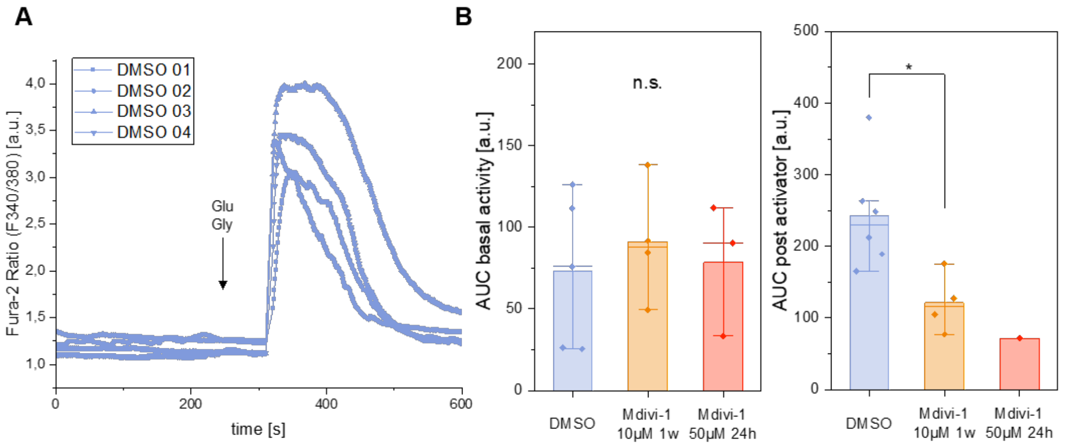


**Figure S 7: Neuronal response to stimuli is reduced Mdivi-1 treated cells.** (A) Representative Fura-2 traces of DMSO treated neurons after addition of glutamate/glycine (100 µM each). (B) Calcium imaging of neurons with long-term Mdivi-1 treatment shows no changes in basal activity (N=2, n=5, n=4, n=4, ANOVA) and less activity after stimulation with activator (N=2, n=6, n=4, n=1, ANOVA).
